# H3K4me1 directs H3K36me2 and H3K36me3 deposition in land plants

**DOI:** 10.1101/2025.06.06.658275

**Authors:** Jiabing Wu, Jiachen Wang, Kangxi Du, Yingping Li, Wenhao Xie, Chenxi He, Qidan Xing, Xiang Li, Xiaoyu Zhu, Zhen Wu, Xiaolong Wu, Wen-Hui Shen, Fei Lan, Jianhua Gan, Bing Liu, Aiwu Dong

**Affiliations:** State Key Laboratory of Genetics and Development of Complex Phenotypes, Department of Biochemistry and Biophysics, Institute of Plant Biology, School of Life Sciences, Fudan University, Shanghai 200438, P.R. China; Liver Cancer Institute, Zhongshan Hospital, Key Laboratory of Carcinogenesis and Cancer Invasion, Ministry of Education, Key Laboratory of Epigenetics and Metabolism, Ministry of Science and Technology, and Institutes of Biomedical Sciences, Fudan University, Shanghai 200438, P.R. China; Institut de Biologie Moléculaire des Plantes, CNRS, Université de Strasbourg, 12 rue du Général Zimmer, 67084 Strasbourg Cédex, France; Shanghai Sci-Tech Inno Center for Infection & Immunity, State Key Laboratory of Genetics and Development of Complex Phenotypes, Collaborative Innovation Center of Genetics and Development, Department of Biochemistry and Biophysics, Fudan University, Shanghai 200438, P.R. China; Ministry of Education Key Laboratory for Biodiversity Science and Ecological Engineering, Institute of Biodiversity Science, School of Life Sciences, Fudan University, Shanghai 200438, P.R. China; State Key Laboratory of Crop Gene Exploration and Utilization in Southwest China, Rice Research Institute, Sichuan Agricultural University, Chengdu 611100, P.R. China; School of Medicine, Chemistry and Chemical Engineering, Taizhou University, 93 East Jichuan Road, Hailing District, Taizhou, Jiangsu 225300, P.R. China

## Abstract

Monomethylation of histone H3 lysine 4 (H3K4me1) marks enhancers in mammals. However, the function of H3K4me1 in plants remains largely unclear. Here, we analyzed the genome-wide distribution of H3K4me1 in diverse species across evolution, revealing a distinctive H3K4me1 distribution pattern in land plants. To explore the function of H3K4me1 in plants, we identified an H3K4me1-specific reader protein, Early heading date 3 (Ehd3), and solved the structure of Ehd3 in complex with the H3K4me1 peptide, revealing a unique binding module differing from the previously reported PHD finger proteins. We further identified an Ehd3-binding protein, SET domain group 724 (SDG724), and the deletion of either *Ehd3* or *SDG724* caused similar defects in plant phenotype and changes in transcriptome and epigenome profiles. Both Ehd3 and SDG724 are enriched at chromatin regions marked by H3K4me1 but not H3K4me2 or H3K4me3. Ehd3 activates the H3K36 methyltransferase SDG724, and H3K36me2/me3 are colocalized with H3K4me1 in the genomes of land plants. Collectively, our results reveal that H3K4me1 directs the establishment of H3K36me2 and H3K36me3 in land plants.

## Main

Histone post-translational modifications (PTMs) contribute to transcriptional regulation and play essential roles in diverse biological processes, including development and response to environmental cues^1, 2^. These modifications, including methylation, acetylation, phosphorylation, and ubiquitination, are dynamically established and removed by specific writer and eraser enzymes, respectively, and are recognized by distinct reader proteins^3, 4^. Methylation of histone H3 lysine 4 (H3K4) is among the most extensively studied PTMs that confers active or repressive transcription depending on its status (mono-, di-, or trimethylation)^5^. H3K4 trimethylation (H3K4me3) is distributed around the transcription start site (TSS) and is associated with gene activation in yeast^6^, animals^7, 8^, and plants^9^. H3K4 dimethylation (H3K4me2) is diversely distributed in eukaryotes: it is enriched at the TSS and associated with gene activation in yeast^6^ and humans^7, 8^, whereas it is located in the gene body region and linked to gene repression in plants^9^. H3K4 monomethylation (H3K4me1) is enriched over the gene body in yeast^6^ and plants^9^, but tends to be concentrated around the TSS in mammals^7, 8^. In addition, H3K4me1 is associated with other PTMs that mark distinct states of enhancers in mammals. Active enhancers are enriched with H3K4me1 and histone H3 lysine 27 acetylation (H3K27ac)^10^. During the intermediate state, primed enhancers primarily exhibit enrichment of H3K4me1^11^. Poised enhancers in pluripotent cells are marked by H3K4me1 and H3K27me3^12^. However, the deposition of H3K4me1 at enhancers is not conserved in plants^13^ and the role of H3K4me1 in plants has yet to be elucidated.

Reader proteins possess specific domains that recognize H3K4 methylation, including the plant homeodomain (PHD), double chromodomain, tandem Tudor domain, and zinc finger CW^3^. To date, a number of structures of PHD finger– H3K4me2/me3 complexes have been determined in animals and plants, whereas the PHD finger in complex with H3K4me1 has not been reported in plants and exclusive readers for H3K4me1 are lacking in animals^14^. The structures of CW or Tudor domains complexed with H3K4me1 have been reported in plants, including the CW domain of SET DOMAIN GROUP 8 (SDG8)^15^ and the Tudor domain of RNA-directed DNA methylation 15 (RDM15)^16^ in Arabidopsis. Nevertheless, the biological function of H3K4me1 decoding in plants remains largely uncharacterized.

Here, we identified a distinct distribution pattern of H3K4me1 in land plants that differs from that observed in yeast and animals. A newly identified PHD finger domain-containing protein, Early heading date 3 (Ehd3), specifically recognizes H3K4me1 and recruits the H3K36 methyltransferase SDG724 to facilitate H3K36me2/me3 deposition. The colocalization of H3K4me1 and H3K36me2/me3 is conserved in land plants, revealing a specific role of H3K4me1 in directing H3K36me2/me3 deposition in plants.

### H3K4me1 exhibits a distinct genome-wide profile in land plants

To determine the function of H3K4me1 in plants, we analyzed the genome-wide distributions of H3K4me1/me2/me3, and the correlation between H3K4 methylations and gene transcription by conducting chromatin immunoprecipitation followed by sequencing (ChIP-seq) and RNA-sequencing (RNA-seq) assays in a variety of plant species, comprising the unicellular aquatic *Chlamydomonas reinhardtii*, the early land plant *Physcomitrium patens*, and the flowering plants *Arabidopsis thaliana* and *Oryza sativa*. By integrating previously published ChIP-seq and RNA-seq data for the unicellular yeast *Saccharomyces cerevisiae*, the invertebrate *Drosophila melanogaster*, and the vertebrates *Mus musculus* and *Homo sapiens*^17–20^, we analyzed the genome-wide H3K4me1/me2/me3 profiles across diverse organisms.

We observed that H3K4me1 exhibited particularly distinct characteristics across the examined species. In animals, comprising *D. melanogaster*, *M. musculus*, and *H. sapiens*, H3K4me1 was enriched near the TSS and gradually decreased along the gene body. In contrast, H3K4me1 was mainly enriched within the gene body in *C. reinhardtii*, *P. patens*, *A. thaliana*, and *O. sativa* (Fig. 1a, b). In yeast and animals, a positive correlation was observed between H3K4me1 and gene transcription (Fig. 1c). In *C. reinhardtii*, a weak correlation between H3K4me1 and gene transcription was detected. In contrast, the land plants *P. patens*, *A. thaliana*, and *O. sativa* displayed a distinctive relationship between H3K4me1 and gene transcription: a high percentage of genes displayed a positive correlation between H3K4me1 and gene transcription, but the most highly expressed genes exhibited lower levels of H3K4me1 (Fig. 1c). Thus, land plants displayed a unique genome-wide distribution pattern for H3K4me1, and a distinct correlation between H3K4me1 and gene transcription.

**Fig. 1:**
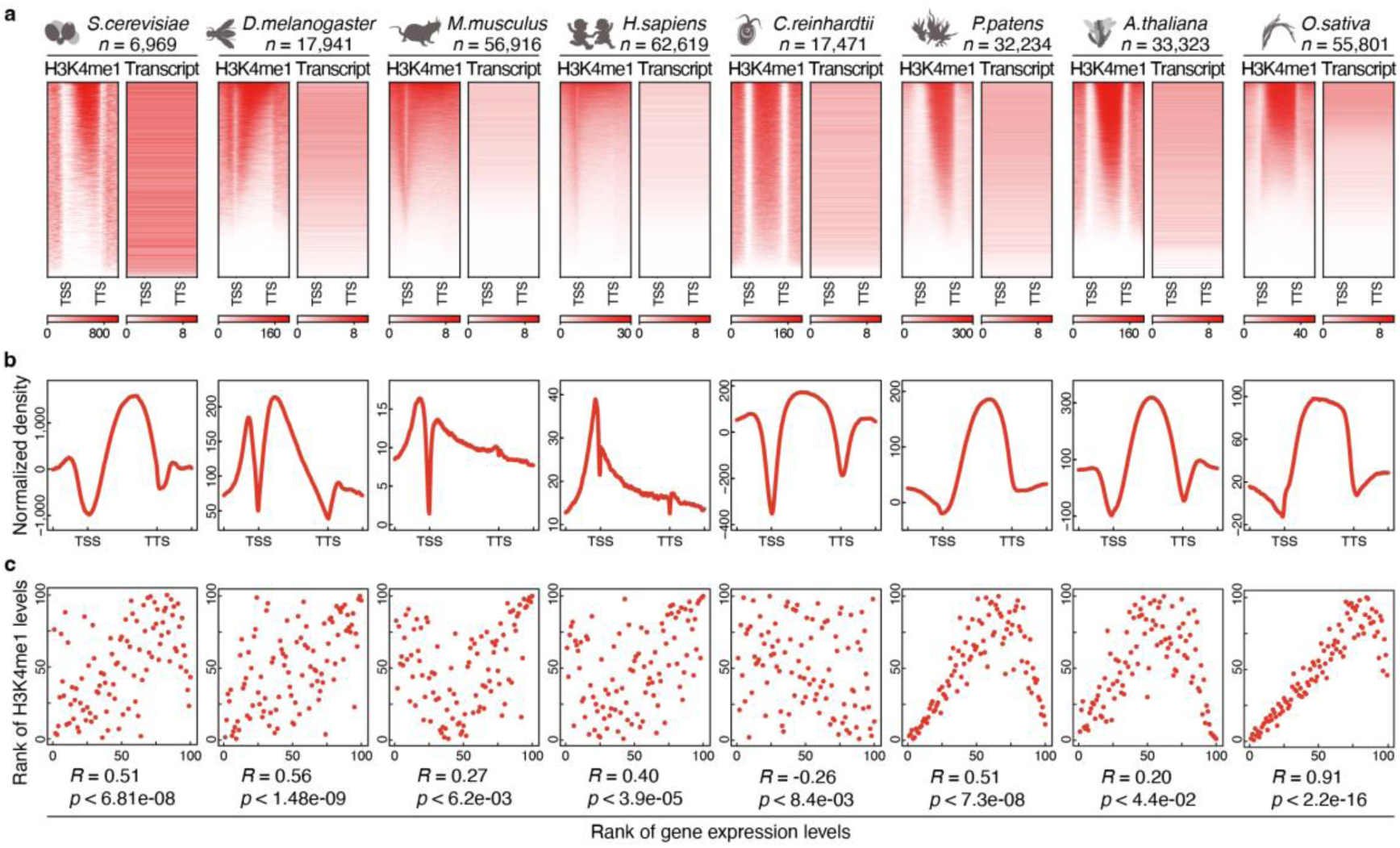
Characterization of the genome-wide distribution of H3K4me1 across diverse species. **a**, Heatmaps showing the individual gene distribution of H3K4me1 (bins per million mapped reads; BPM), together with their associated transcription level (transcripts per million; TPM). All genes are sorted based on enrichment of H3K4me1. **b**, Integrative genomic distribution of H3K4me1. **c**, Scatter plots showing the correlation between H3K4me1 level and gene transcription level. The species (from left to right) are *Saccharomyces cerevisiae*, *Drosophila melanogaster*, *Mus musculus*, *Homo sapiens*, *Chlamydomonas reinhardtii*, *Physcomitrium patens*, *Arabidopsis thaliana*, and *Oryza sativa*, respectively. The transcribed genes enriched with the H3K4me1 modification were divided into 100 groups based on the levels of their transcription and modification. The average values of each group of modifications and transcription were calculated for analysis. The heatmaps and plots present the region from 3 kb upstream of the transcription start site (TSS) to 3 kb downstream of the transcription termination site (TTS). For scatter plots, Spearman’s rank correlation coefficient (*R*) indicates the correlation between the methylation level and gene transcription level. The *p*-value was determined based on Spearman’s rank correlation test.

Unlike animals, in which two peaks around the TSS were observed, H3K4me2 was distributed over the entire gene body in land plants (Extended Data Fig. 1a, b). Consistent with a previous study, a positive correlation between H3K4me2 level and gene transcription level was observed in yeast and animals (Extended Data Fig. 1c). However, H3K4me2 was negatively correlated with gene transcription in land plants, consistent with our previous report^9^ (Extended Data Fig. 1c). The H3K4me3 distributions in all investigated species were similar, peaking close to the TSS (Extended Data Fig. 1d, e). Moreover, H3K4me3 consistently displayed a positive correlation with gene transcription in all species examined, highlighting the conserved association of H3K4me3 with gene activation in eukaryotes (Extended Data Fig. 1f).

### Ehd3, a PHD finger domain-containing protein, specifically recognizes H3K4me1

The unique distribution pattern and distinct correlation with gene transcription suggested that H3K4me1 may play a unique role in land plants. Given that the epigenetic information encoded by histone PTMs are interpreted by specialized reader proteins, we thus investigated the function of H3K4me1 in land plants by searching for its specific reader proteins. Given the distinct distribution of H3K4me1 between *C. reinhardtii* and *P. patens*, we screened the PHD finger domain-containing proteins, which represent the largest family of H3K4 reader proteins in plants, aiming to discover potential specific reader proteins conserved in land plants but not in *C. reinhardtii*. We determined that a PHD finger domain-containing protein (A0A2K3D188) in *C. reinhardtii* has a different domain structure compared with its homologs in *P. patens*, *A. thaliana*, and *O. sativa*, which have evolved tandem PHD finger domains (Fig. 2a). Ehd3, the rice homolog of A0A2K3D188, was selected for in-depth study (Fig. 2b). To explore the histone modification decoding mode of Ehd3, we purified the full-length recombinant Ehd3 protein and performed an *in vitro* binding assay using a histone peptide microarray with diverse histone PTMs. Ehd3 exhibited a strong preference for binding to H3K4me1 over other modifications (Extended Data Fig. 2a). A biotinylated histone peptide pull-down assay similarly showed that Ehd3 interacted exclusively with the H3K4me1 peptide, but not with other tested modifications, including H3K4me0, H3K4me2, and H3K4me3 (Fig. 2c and Supplementary Table 1). Further investigation showed that individual PHD1, PHD2, or PHD3 failed to bind to H3K4me1, whereas the tandem PHD finger domain (PHD2–PHD3) was essential and sufficient for recognition of H3K4me1 (Fig. 2d). Isothermal titration calorimetry (ITC) analysis further confirmed robust interaction between Ehd3 and the H3K4me1 peptide with a dissociation constant (*K*_d_) of 1.7 μM (Fig. 2e and Supplementary Tables 1–2). In summary, Ehd3 specifically recognized H3K4me1 through its tandem PHD finger domain.

**Fig. 2:**
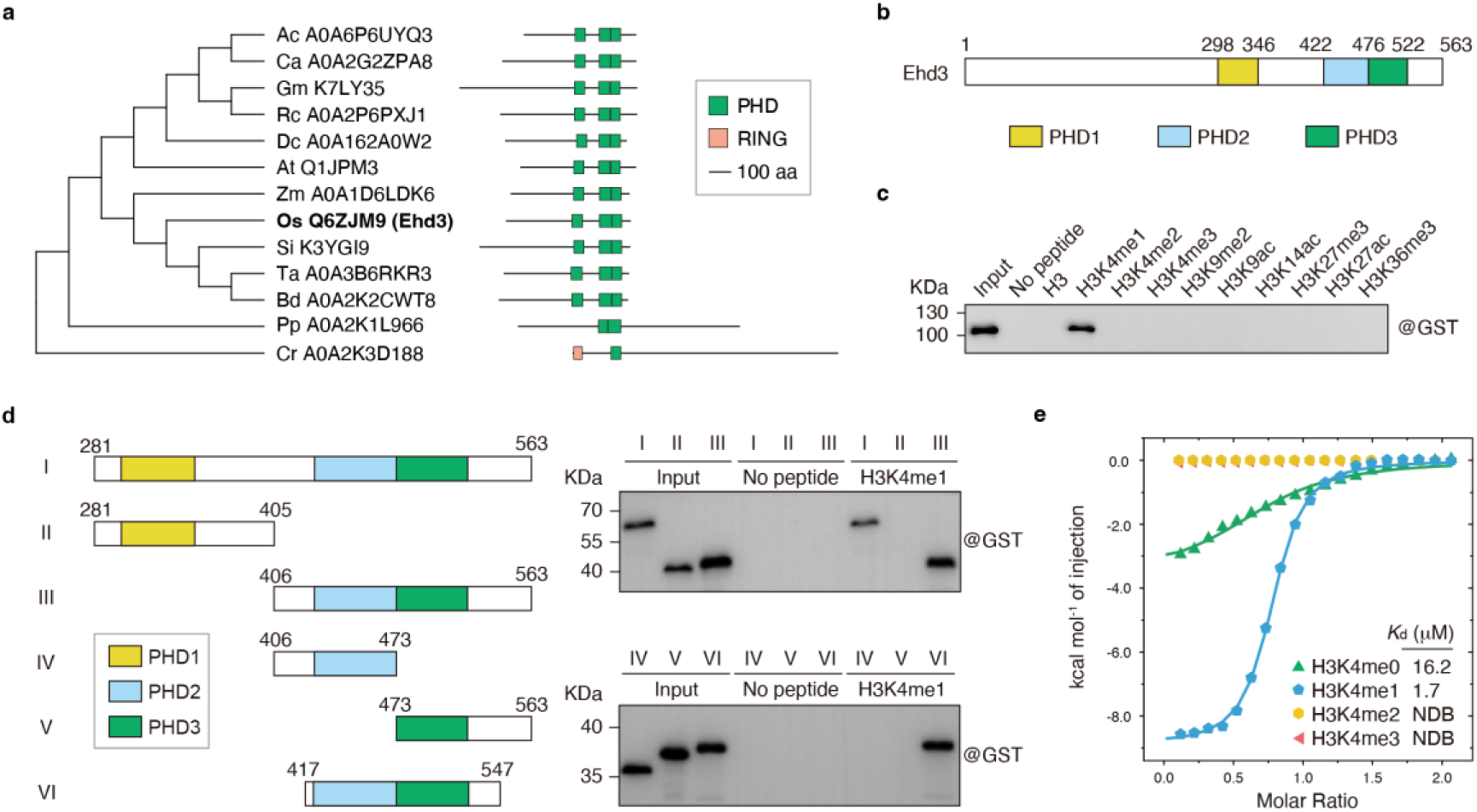
Ehd3 specifically recognizes H3K4me1. **a**, Neighbor-joining dendrogram (left) and domain architecture (right) of Ehd3 homologous proteins in plants. The neighbor-joining dendrogram was constructed based on the tandem plant homeodomain (PHD) amino acid sequence alignment. Ac, *Coffea arabica*; Ca, *Capsicum annuum*; Gm, *Glycine max*; Rc, *Rosa chinensis*; Dc, *Daucus carota*; At, *Arabidopsis thaliana*; Zm, *Zea mays*; Os, *Oryza sativa*; Si, *Setaria italica*; Ta, *Triticum aestivum*; Bd, *Brachypodium distachyon*; Pp, *Physcomitrium patens*; Cr, *Chlamydomonas reinhardtii*. UniProt (https://www.uniprot.org) entries for each protein are given after the species abbreviation. **b**, Schematic representation of the full-length Ehd3 protein. The individual PHD finger domains are indicated by different colored boxes. **c**, Immunoblotting analysis of histone peptide pull-down using the full-length Ehd3 fused with glutathione-*S*-transferase (GST). **d**, Schematic representation of different truncated Ehd3 proteins fused with GST (left) and immunoblotting analysis of H3K4me1-peptide pull-down (right). Roman numerals indicate the corresponding truncations. **e**, Isothermal titration calorimetry results showing the binding affinities of the full-length Ehd3 with different H3K4 methylated peptides. NDB, no detectable binding.

### Structure of Ehd3 tandem PHD finger domain in complex with H3K4me1 peptide

To clarify the molecular mechanism underlying Ehd3 recognition of H3K4me1, we solved the crystal structure of Ehd3_417–547_ bound to a H3K4me1 peptide (A_1_RT-Kme1-QTARKSTG) at atomic resolution (1.5 Å, Supplementary Table 3). Ehd3_418–540_ adopts an L-shaped overall architecture and the peptide binds across an acidic groove formed by the PHD2 and PHD3 domains (Fig. 3a). Both PHD2 and PHD3 of Ehd3 adopt interleaved topologies chelating two Zn^2+^ ions (Extended Data Fig. 2b, c), while the H3K4me1 peptide adopts an extended conformation (Extended Data Fig. 2d, e). Partial atoms of Gln5, Arg8, and Lys9 are invisible in the map, suggesting flexible disposition of these residues (Extended Data Fig. 2d, e). The side chains of Ala1 and Arg2 of the H3K4me1 peptide reside in two surface channels separated by Asp493 of Ehd3. The monomethylated Lys4 inserts into a deep pocket adjacent to the Arg2-binding channel, forming a ‘Y’-shaped conformation with Ala1 and Arg2 that anchor the binding interface (Fig. 3b). The main-chain N atom of Ala1 forms two stable hydrogen bond (H-bond) interactions with the main-chain O atoms of Pro512 and Gly514, respectively, whereas the methyl group of Ala1 forms a hydrophobic interaction with the aromatic ring of Pro512 (Extended Data Fig. 2f). The main-chain N and O atoms of Arg2 form two water-mediated H-bonds and one direct H-bond interaction with Met491 and Asp493, respectively. Via the guanidyl group, the side chain of Arg2 forms three stable H-bond interactions with Tyr446, Cys492, and Asp496 (Fig. 3c and Extended Data Fig. 2g, h).

**Fig. 3:**
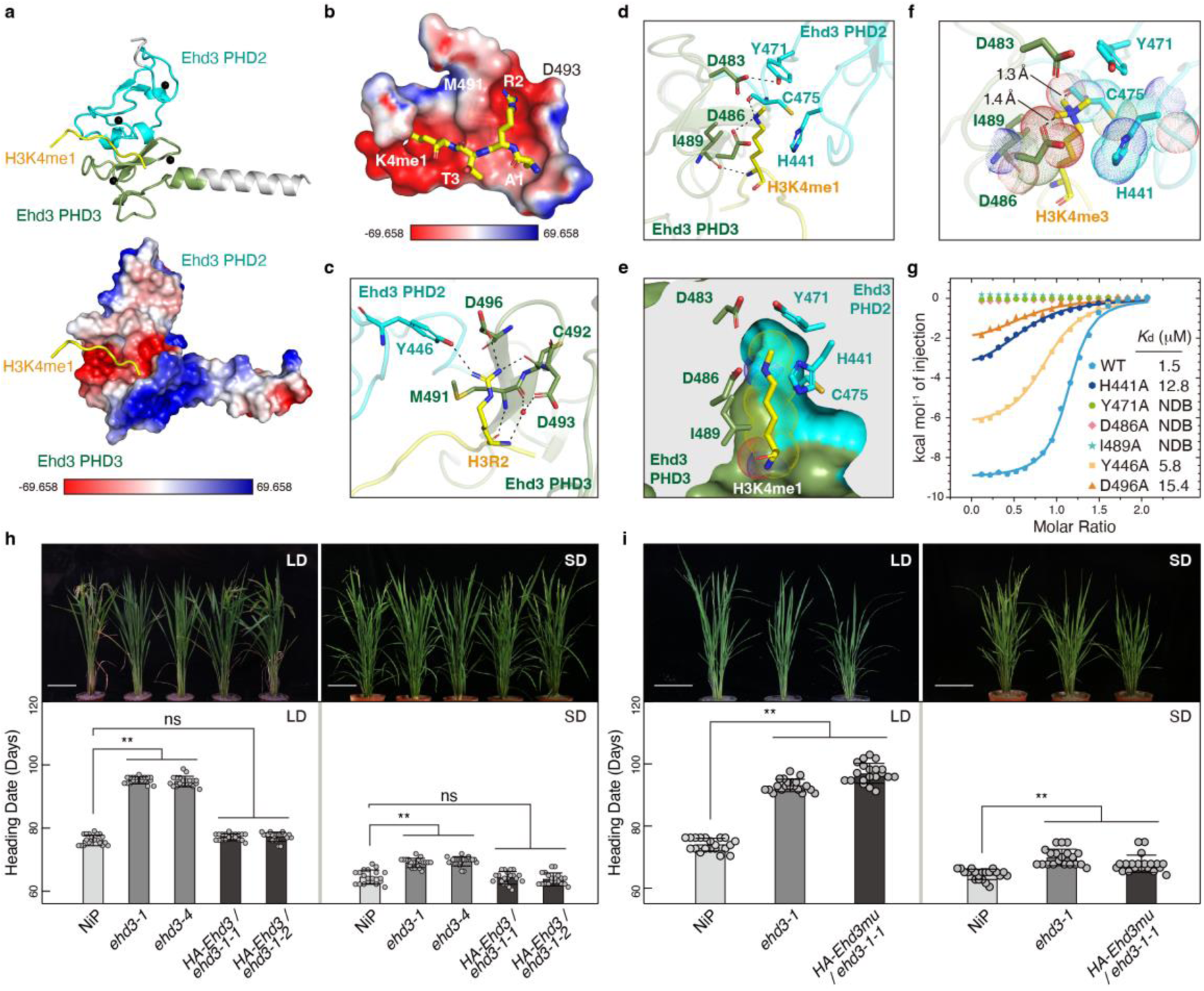
Crystal structure of Ehd3 in complex with H3K4me1 peptide and function of Ehd3 in plant development. **a**, Overall structure (upper panel) and electrostatic surface view (lower panel) of the Ehd3 tandem plant homeodomain (PHD) finger domain in complex with the H3K4me1 peptide. PHD2 and PHD3 of the Ehd3 tandem PHD finger domain are shown in cyan and green, respectively, and the H3K4me1 peptide is in yellow. Blue and red shades on the surface indicate positive and negative charges, respectively. **b**, Electrostatic surface showing the details of the Ehd3 tandem PHD finger domain–H3K4me1 complex. **c**, Detailed interaction between the Ehd3 tandem PHD finger domain and H3R2. Hydrogen bonds are indicated by dashed lines. **d**, Detailed interaction of the Ehd3 tandem PHD finger domain and H3K4me1. Hydrogen bonds are indicated by dashed lines. **e**, Ehd3 tandem PHD finger domain cut-away view of H3K4me1 recognition. **f**, Simulated interactions between the Ehd3 tandem PHD finger domain and H3K4me3. **g**, Isothermal titration calorimetry results showing the binding affinities between the wild-type or site-mutated Ehd3 tandem PHD finger domain with the H3K4me1 peptide. NDB, no detectable binding. **h**, Representative image (upper panel) and heading date (lower panel) of wild-type rice ‘Nipponbare’ (NiP), *ehd3-1*, *ehd3-4*, *HA-Ehd3/ehd3-1-1*, and *HA-Ehd3/ehd3-1-2* plants grown under long-day (LD) and short-day (SD) photoperiods. **i**, Representative image (upper panel) and heading date (lower panel) of wild-type NiP, *ehd3-1*, and *HA-Ehd3mu/ehd3-1-1* plants grown under LD and SD photoperiods. Scale bar, 20 cm. Values are the mean ± standard deviation of 20 individual plants. Statistical significance was determined with one-way ANOVA: ** *p*-value < 0.01; ns (not significant).

In contrast to trimethylated Lys4-binding pockets present in many PHD proteins, which are composed of several conserved aromatic residues within a single PHD domain (Extended Data Fig. 3), the monomethylated Lys4-binding pocket of Ehd3 is formed by residues from both PHD2 (His441, Tyr471, and Cys475) and PHD3 (Asp483, Asp486, and Ile489) (Fig. 3d and Extended Data Fig. 2h, i). The monomethylated Lys4 NZ forms H-bond interactions with the main-chain O atom of PHD2 Cys475 and the side-chain OD1 of PHD3 Asp486. The methyl group and the CE atom of Lys4 hydrophobically interact with the side chains of PHD2 Tyr471 and His441, respectively. The main chains of Lys4 and PHD3 Ile489 form a H-bond and the side chain of Ile489 hydrophobically interacts with CB and CD of Lys4. The OH group of PHD2 Tyr471 forms a H-bond interaction with the OD2 of PHD3 Asp483 to stabilize the binding pocket (Fig. 3d and Extended Data Fig. 2h). As a result, the side chain of monomethylated Lys4 perpendicularly and compactly inserts into the pocket, which harbors a hydrophobic environment created by PHD2 and a polar environment created by PHD3 (Fig. 3e). Our modeling study suggests that the introduction of a second or third methyl group to Lys4 will lead to a clash with Cys475 and/or Asp486, which will prevent the binding of Ehd3 to H3K4me2 or H3K4me3, implying the H3K4me1 specificity of Ehd3 (Fig. 3f). In addition to the monomethylated Lys4, the side chain of PHD2 His441 NE forms one stable H-bond interaction with the main-chain O atom of Gln5 of the peptide. Residues 6–9 of the peptide mainly form direct or water-mediated H-bond interactions with symmetry-related Ehd3 molecules, whereas residues 10–12 of the peptide are completely disordered in the structure (Extended Data Fig. 2h, j, k). In agreement with the structural observations, the Ala substitution for each H3K4me1 binding-pocket residue markedly reduced the affinity of the Ehd3 tandem PHD finger for the H3K4me1 peptide (Fig. 3g and Supplementary Table 2). The dissociation constant of the wild-type Ehd3 tandem PHD to H3K4me1 peptide was comparable to that of the full-length Ehd3 (Figs. 2e, 3g), excluding the potential contribution of PHD1 to specific recognition of H3K4me1.

Given that Double PHD Finger 3b (DPF3b) is a well-studied epigenetic factor that recognizes histone modifications through a tandem PHD finger domain^21^, we generated a sequence alignment of the tandem PHD of Ehd3 and DPF3b. Most H3K4me1-binding sites in Ehd3 were not conserved in DPF3b (Extended Data Fig. 3a). The tandem PHD finger domain in Ehd3 forms a narrow and sealed pocket, whereas DPF3b relies on a single PHD finger domain to create an open pocket for H3K14ac binding (Fig. 3e and Extended Data Fig. 3c, d). In addition, we compared the reported structures of the H3K4me3-binding PHD finger domains in the Bromodomain PHD finger transcription factor (BPTF)^22^, PHD finger protein 2 (PHF2)^23^, YNG1^24^, and Inhibitor of growth protein 2 (ING2)^25^ with that of Ehd3–H3K4me1. The interaction between the PHD finger domain and H3K4me3 is highly conserved in these proteins and the critical residues are not conserved in the three PHD finger domains of Ehd3 (Extended Data Fig. 3b, e–h). In most cases, a single PHD finger domain is sufficient for recognition of histone modifications, highlighting a novel binding mode of the tandem PHD finger domain within Ehd3 to specifically recognize H3K4me1.

### Ehd3–H3K4me1 interaction is required for proper timing of flowering in rice

To further investigate the functional significance of Ehd3–H3K4me1 binding *in vivo*, we first generated CRISPR/Cas9 mutants targeting *Ehd3* in the ‘Nipponbare’ (NiP) background. Two single guide RNAs (sgRNAs) targeting the second exon of *Ehd3* were designed. We selected mutants with two independent transgene-free mutations, one with a 1-bp deletion at target site 1 and a 9-bp deletion at target site 2 (*ehd3-1*), and the other with a 4-bp deletion at target site 1 and a 1-bp insertion at target site 2 (*ehd3-4*), for further analysis (Extended Data Fig. 4a). The *ehd3-1* and *ehd3-4* mutant plants displayed a delayed-flowering phenotype under both natural long-day (LD) and short-day (SD) photoperiods (Extended Data Fig. 4b), consistent with the reported phenotypes of *ehd3* mutants in the *O. sativa* subsp. *japonica* ‘Tohoku IL9’ background^26, 27^. We then generated the plants *HA-Ehd3/ehd3-1-1* and *HA-Ehd3/ehd3-1-2*, expressing *Ehd3* fused with the *4×HA* tag driven by its native promoter in the *ehd3-1* mutant background (Extended Data Fig. 4c). The delayed-flowering phenotype of *ehd3-1* was fully rescued upon the expression of *HA-Ehd3* (Fig. 3h), indicating that the delayed-flowering phenotype is attributable to mutations in *Ehd3*.

In addition, we generated transgenic plants expressing a mutated version of *Ehd3* by substituting two critical Ehd3–H3K4me1-interacting residues, Tyr471 and Asp496, with Ala under the control of its native promoter in the *ehd3-1* mutant background (Extended Data Fig. 4c). The transgenic line (*HA-Ehd3mu/ehd3-1-1*) carrying these site-specific mutations exhibited a delayed-flowering phenotype similar to that of the *ehd3-1* mutant (Fig. 3i), suggesting that the mutation of critical residues responsible for Ehd3–H3K4me1 interaction within Ehd3 failed to rescue the phenotype of the *ehd3-1* mutant. All of the aforementioned findings showed that binding to H3K4me1 is critical for the biological function of Ehd3 in rice.

### Ehd3 physically interacts with the H3K36 methyltransferase SDG724

To further investigate the role of H3K4me1 through Ehd3 in plants, we identified the co-factors of Ehd3 by immunoprecipitation followed by mass spectrometry (IP-MS). Transgenic plants were generated harboring *Enhanced Green Fluorescent Protein* (*EGFP*) fused to *Ehd3* driven by the rice *Ubiquitin* promoter (Extended Data Fig. 4d). Among the candidates for Ehd3-binding proteins identified in the IP-MS experiment, the H3K36-specific methyltransferase SDG724 was identified^28^ (Fig. 4a and Supplementary Table 4). Therefore, transgenic plants expressing *Yellow Fluorescent Protein* (*YFP*) fused to *SDG724* driven by the *Ubiquitin* promoter were generated (Extended Data Fig. 4e). Ehd3 was among the SDG724-binding candidates identified by IP-MS (Fig. 4a and Supplementary Table 5), suggesting the possible interaction between Ehd3 and SDG724. Yeast two-hybrid, glutathione-*S*-transferase (GST) pull-down assays, co-immunoprecipitation, and bimolecular fluorescence complementation (Fig. 4b–e and Extended Data Fig. 4f) assays consistently confirmed the physical interaction between Ehd3 and SDG724 in rice.

**Fig. 4:**
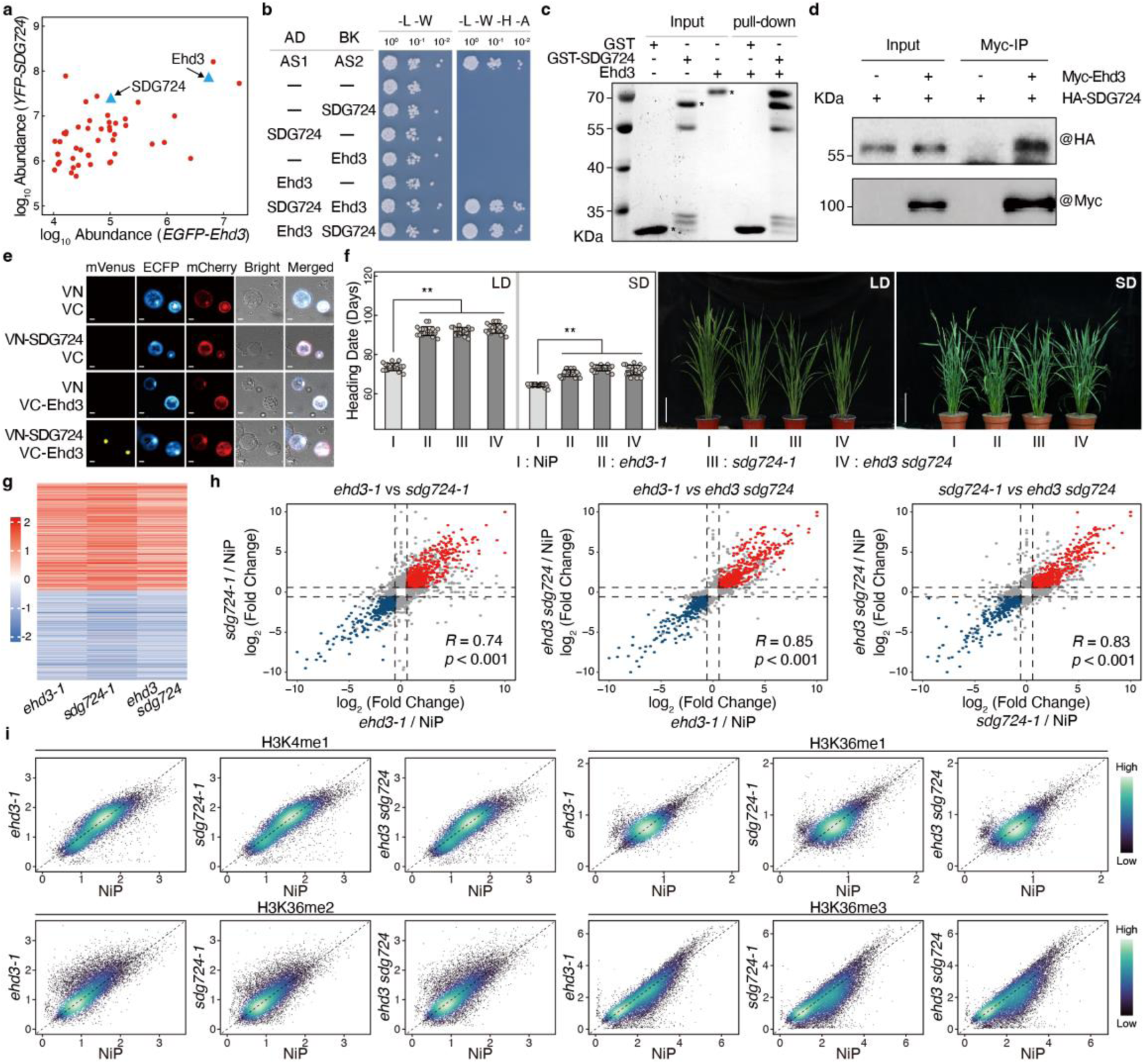
Ehd3 cooperates with SDG724 to regulate gene transcription and H3K36 methylation. **a**, Scatter plot representing mass spectrometry results, displaying the protein abundance identified in *EGFP-Ehd3* and *YFP-SDG724* immunoprecipitation analyses. The *x*-axis shows the log_10_-abundance of proteins in the *EGFP-Ehd3* IP-MS assay, and the *y*-axis shows the log_10_-abundance of proteins in the *YFP-SDG724* IP-MS assay. Ehd3 and SDG724 are marked with blue triangles. **b**, Yeast two-hybrid assay showing the interaction between Ehd3 and SDG724 in yeast cells. AS1 and AS2 were used as the positive control. **c**, *In vitro* glutathione-*S*-transferase (GST) pull-down assay showing the interaction between Ehd3 and SDG724. Ehd3 was pulled down by GST-SDG724 but not GST. GST, GST-SDG724, and Ehd3 are marked by asterisks in the gel. **d**, Co-immunoprecipitation assay showing the interaction of Ehd3 and SDG724 *in vivo*. Proteins from rice protoplasts expressing *Myc-Ehd3* and *HA-SDG724* were extracted. Myc-Ehd3 and HA-SDG724 were detected by western blotting using anti-Myc and anti-HA antibodies, respectively. **e**, Bimolecular fluorescent complementation assay showing the interaction of Ehd3 and SDG724 in rice protoplasts. Scale bar, 10 μm. **f**, Heading date (left) and representative images (right) of wild-type rice ‘Nipponbare’ (NiP), *ehd3-1*, *sdg724-1*, and *ehd3 sdg724* plants grown under long-day (LD) and short-day (SD) photoperiods. Scale bar, 20 cm. Values are the mean ± standard deviation of 20 individual plants. Statistical significance was determined with one-way ANOVA: ** *p*-value < 0.01. **g**, Heatmap showing the log_2_(fold change) of differentially expressed genes (DEGs) in the *ehd3-1*, *sdg724-1*, and *ehd3 sd724* mutants compared with the wild-type NiP. Up- and down-regulated genes are indicated in red and blue, respectively. **h**, Scatter plots showing the highly correlated DEGs (*p*-value ≤ 0.05 and |fold change| ≥ 1.5) in the *ehd3-1*, *sdg724-1*, and *ehd3 sdg724* mutants. **i**, Scatter plots comparing H3K4me1, H3K36me1, H3K36me2, and H3K36me3 levels between the wild-type NiP and the *ehd3-1*, *sdg724-1*, and *ehd3 sdg724* mutants. Each dot represents the histone methylation level of one peak corrected with MAnorm. The transition from deep blue to cyan indicates the increase in dot density.

### Deletion of *Ehd3* or *SDG724* leads to similar changes in plant phenotype, transcriptome, and epigenome

Both Ehd3 and SDG724 promote flowering under SD and LD photoperiods^28^ (Fig. 4f). To investigate the genetic relationship between *Ehd3* and *SDG724*, we crossed the *ehd3-1* and *sdg724-1* mutants to generate the *ehd3 sdg724* double mutant. The *ehd3 sdg724* double mutant displayed a similar late-flowering phenotype to that of each single mutant under both SD and LD photoperiods (Fig. 4f), implying that Ehd3 and SDG724 promote flowering through the same genetic pathway.

To evaluate how Ehd3 and SDG724 regulate gene transcription at the genome scale, we conducted RNA-seq analyses of the wild-type NiP and the *ehd3-1*, *sdg724-1*, and *ehd3 sdg724* mutants. Hundreds of differentially expressed genes (DEGs) were identified in the *ehd3-1*, *sdg724-1*, and *ehd3 sdg724* mutants compared with NiP (Extended Data Fig. 5a and Supplementary Table 6). Significant overlap in the up- and down-regulated genes among the single and double mutants was observed (Extended Data Fig. 5b, c). The overlapping DEGs displayed similar transcription changes in the *ehd3-1*, *sdg724-1*, and *ehd3 sdg724* mutants (Fig. 4g). Furthermore, pairwise scatterplot analyses indicated a strong correlation in the DEGs among the *ehd3-1*, *sdg724-1*, and *ehd3 sdg724* mutants (Fig. 4h), implying cooperative regulation of gene transcription by Ehd3 and SDG724.

In addition, we investigated the global distributions of H3K4me1/me2/me3, H3K36me1/me2/me3, and H3 in NiP and the *ehd3-1*, *sdg724-1*, and *ehd3 sdg724* mutants via ChIP-seq analysis. Gene-by-gene scatterplots and heatmaps revealed no alterations in the distributions of H3K4me2/me3, subtle changes for H3K4me1 and H3K36me1, and significant changes in H3K36me2/me3 distributions in the *ehd3-1*, *sdg724-1*, and *ehd3 sdg724* mutants compared with those of NiP (Fig. 4i and Extended Data Fig. 6a, b). As a control, the H3 distribution in these mutants was comparable to that in NiP (Extended Data Fig. 6c). Therefore, Ehd3 cooperates with SDG724 to regulate H3K36me2/me3 distributions, gene transcription, and plant development.

### SDG724 collaborates with Ehd3 to deposit H3K36me2/me3 at genes marked by H3K4me1

The genome-wide occupancy of Ehd3 was analyzed using the *HA-Ehd3/ehd3-1-1* transgenic plants. Ehd3 was preferentially enriched at genes marked with H3K4me1, but not H3K4me2 or H3K4me3, compared with the random control genes, which supported the hypothesis that Ehd3 specifically recognizes H3K4me1 *in vivo* (Fig. 5a, b, Extended Data Fig. 6d, and Supplementary Tables 7–8). A ChIP-seq assay to assess the global distribution of SDG724 in the *YFP*-*SDG724* transgenic plants showed that SDG724 exhibited a notable co-occupancy with Ehd3 at H3K4me1-enriched genes (Fig. 5a, b, Extended Data Fig. 6d, and Supplementary Tables 7–8). Gene-by-gene heatmaps revealed that loss of either *Ehd3* or *SDG724* caused significant changes in H3K36me2/me3 distributions within the genes enriched with H3K4me1 (Fig. 5c). In detail, H3K36me2 occupancy decreased in the 3′ regions, whereas H3K36me3 significantly decreased across the gene body in the *ehd3-1*, *sdg724-1*, and *ehd3 sdg724* mutants compared with NiP (Fig. 5c). Collectively, these results indicated that Ehd3 and SDG724 together bind to genes enriched with H3K4me1 and promote deposition of H3K36me2 and H3K36me3 (Fig. 5d and Extended Data Fig. 6e).

**Fig. 5:**
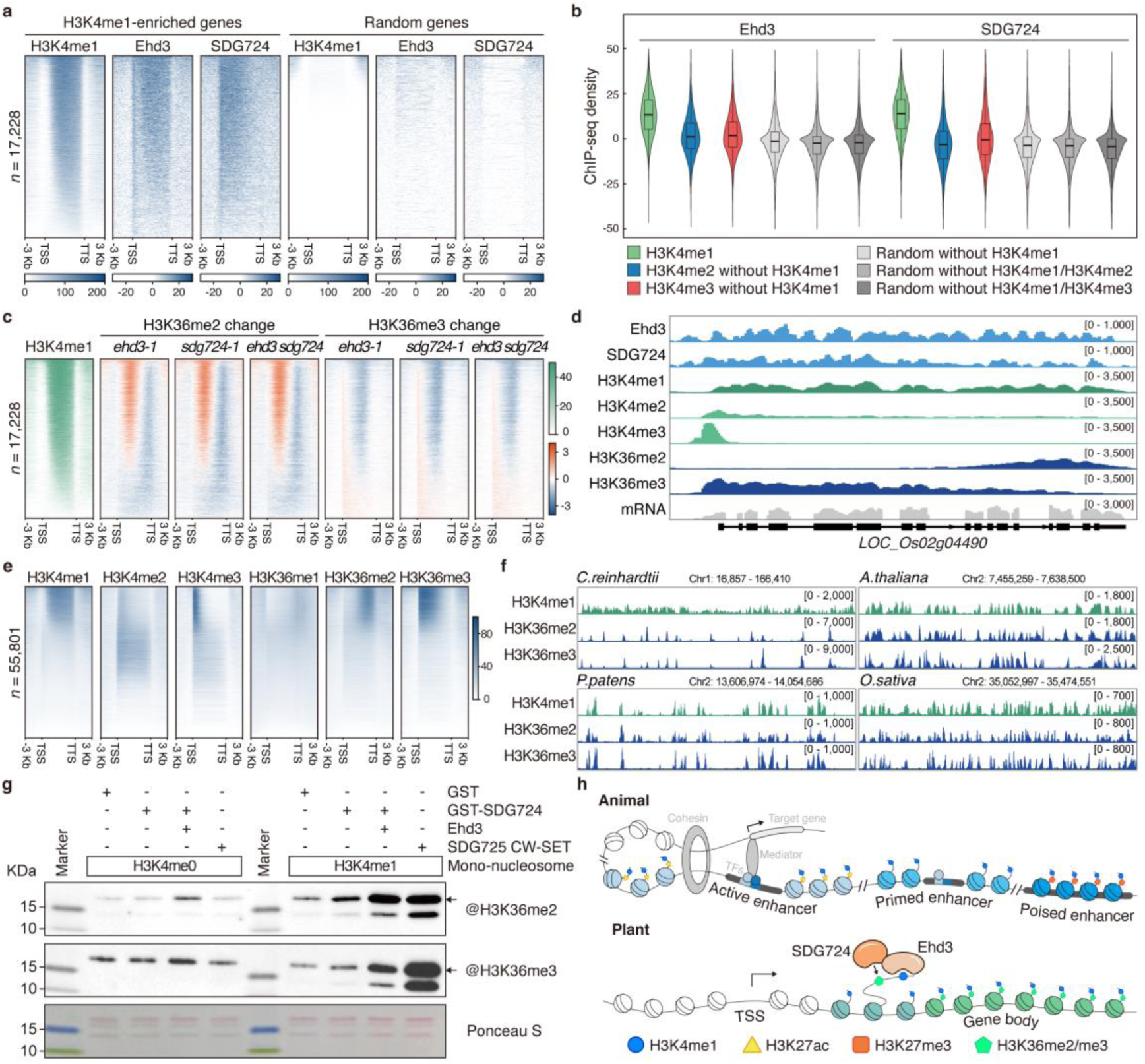
H3K4me1 is specifically colocalized with H3K36me2/me3 in land plants. **a**, Heatmaps showing the enrichment levels (reads per kilobase per million mapped reads; RPKM) of Ehd3 and SDG724 within H3K4me1-enriched genes from 3 kb upstream of the transcription start site (TSS) to 3 kb downstream of the transcription termination site (TTS). An equal number of genes lacking H3K4me1 enrichment were randomly selected as the control. All genes are sorted based on the enrichment of H3K4me1. Numbers of genes are shown on the left. **b**, Violin plots and box plots showing Ehd3 (left) and SDG724 (right) levels within H3K4me1-enriched (green), H3K4me2 without H3K4me1-enriched (blue), or H3K4me3 without H3K4me1-enriched (red) genes. An equal number of genes lacking H3K4me1, H3K4me1/H3K4me2, or H3K4me1/H3K4me3 enrichment were randomly selected as the control. **c**, Heatmaps showing changes in H3K36me2 and H3K36me3 levels (bins per million mapped reads; BPM) within H3K4me1-enriched genes, compared with the wild-type rice ‘Nipponbare’ (NiP), from 3 kb upstream of the TSS to 3 kb downstream of the TTS in the *ehd3-1*, *sdg724-1*, and *ehd3 sdg724* mutants. All genes are sorted based on enrichment of H3K4me1 (BPM) identified in the wild-type NiP. Numbers of genes are shown on the left. **d**, Integrative Genomics Viewer images of ChIP-seq data showing the distributions of Ehd3, SDG724, H3K4me1, H3K4me2, H3K4me3, H3K36me2, and H3K36me3 along the representative gene in the wild-type NiP. **e**, Heatmaps showing the individual gene distribution of H3K4me1/me2/me3 and H3K36me1/me2/me3 (BPM) from 3 kb upstream of the TSS to 3 kb downstream of the TTS in the wild-type NiP. All genes are sorted based on the enrichment of H3K4me1. Numbers of genes are shown on the left. **f**, Integrative Genomics Viewer images of ChIP-seq data showing the H3K4me1-, H3K36me2-, and H3K36me3-enrichment profile plots of the representative region in *Chlamydomonas reinhardtii*, *Physcomitrium patens*, *Arabidopsis thaliana*, and *Oryza sativa*. **g**, Histone methyltransferase assay showing that Ehd3 stimulates the H3K36 methyltransferase activity of SDG724. H3K4me0 and H3K4me1 mononucleosomes were used as the substrates. Glutathione-*S*-transferase (GST) and truncated SDG725 (containing the CW and SET domains) were used as negative and positive controls, respectively. **h**, Role of H3K4me1 in animals and plants. Upper panel: In animals, H3K4me1 marks various enhancer states. Within a given tissue, a subset of poised enhancers is characterized by H3K4me1 and H3K27me3. Prior to activation, enhancers existing in a primed state marked only H3K4me1. Active enhancers, which lack nucleosomes, are adjacent to nucleosomes that exhibit marks of H3K4me1 and H3K27ac. These active enhancers bind transcription factors that target specific genes through looping interactions mediated by cohesion and other protein complexes. Lower panel: In plants, H3K4me1 directs the establishment of H3K36me2/me3. The tandem plant homeodomain (PHD) finger domain-containing protein Ehd3 specifically recognizes H3K4me1 and interacts with the H3K36 methyltransferase SDG724. This interaction stimulates the catalytic activity of SDG724, facilitating the deposition of H3K36me2/me3, which results in the colocalization of H3K4me1 and H3K36me2/me3 in plants.

### H3K4me1 directs H3K36me2 and H3K36me3 deposition in land plants

Correlation analyses showed that H3K4me1 was positively correlated with H3K36me2 (*R* = 0.54) and H3K36me3 (*R* = 0.64), but not with other tested histone modifications, implying that H3K4me1 is associated with H3K36me2/me3 in rice^29, 30^ (Extended Data Fig. 7a). SDG724 is required for deposition of H3K36me2/me3 at H3K4me1-enriched genes, supporting the positive correlation between H3K4me1 and H3K36me2/me3 in rice (Extended Data Fig. 7a). In addition, gene-by-gene heatmaps showed that genes enriched with H3K4me1 exhibited significant enrichment of H3K36me2/me3 (Fig. 5e and Supplementary Tables 7–8), supporting the observation that H3K4me1 specifically colocalizes with H3K36me2/me3 in rice. To investigate if the colocalization of H3K4me1 and H3K36me2/me3 is conserved in plants, we performed ChIP-seq analysis of H3K36me2/me3 in *C. reinhardtii* and *P. patens*, and integrated published H3K36me2/me3 ChIP-seq data for *S. cerevisiae*, *D. melanogaster*, *M. musculus*, *H. sapiens*, and *A. thaliana*^31–33^ (Fig. 5f and Extended Data Fig. 7b). In contrast to the diverse patterns of H3K4me1 and H3K36me2/me3 distributions in *S. cerevisiae*, *C. reinhardtii*, *D. melanogaster*, *M. musculus*, and *H. sapiens*, the distributions of H3K4me1 and H3K36me2/me3 displayed high similarity in *P. patens*, *A. thaliana*, and *O. sativa* (Fig. 5f and Extended Data Fig. 7b). Therefore, a distinct evolutionary trend for colocalization of H3K4me1 and H3K36me2/me3 in land plants was implied.

To investigate the molecular mechanism linking H3K4me1 and H3K36me2/me3, a histone methyltransferase assay was performed using H3K4me0- or H3K4me1-modified mononucleosomes. The truncated rice H3K36 methyltransferase SDG725 containing the CW and SET domains, and GST served as positive and negative controls, respectively. When incubated with H3K4me0 nucleosomes, neither SDG724 nor SDG725 showed obvious methyltransferase activity (Fig. 5g). However, using H3K4me1-modified nucleosomes as the substrate, the SDG724 and SDG725 activities greatly increased. Moreover, the SDG724 activity further increased in the presence of Ehd3, indicating that Ehd3 is required to stimulate the catalytic activity of SDG724 (Fig. 5g). The CW domain of SDG8, a homolog of SDG725 in *A. thaliana*, is responsible for binding to H3K4me1^15^, and the critical residues of SDG8 involved in H3K4me1 binding are conserved in *O. sativa* SDG725 (Extended Data Fig. 7c), supporting the hypothesis that SDG725 is able to recognize H3K4me1 through its CW domain. Collectively, recognition of H3K4me1-modified nucleosomes by the interacting reader protein or internal reading domain is essential for plant H3K36 methyltransferases, leading to H3K36me2/me3 deposition on chromatin. Taken together, these results revealed that H3K4me1 directs H3K36me2/me3 deposition in land plants.

## Discussion

In this study, we identified a specific H3K4me1 reader protein, Ehd3, and revealed a crucial role of H3K4me1 in facilitating H3K36me2/me3 deposition in rice. Ehd3 recognizes H3K4me1 through its tandem PHD finger domain and recruits the H3K36 methyltransferase SDG724 to H3K4me1-enriched chromatin regions. Furthermore, interaction with Ehd3 enhances the catalytic activity of SDG724 in H3K36 methylation on H3K4me1-modified nucleosomes. The association between H3K4me1 and H3K36me2/me3 across the genome is conserved in land plants (Fig. 5h). The crystal structure of the Ehd3 tandem PHD finger domain in complex with H3K4me1 reveals a unique binding mode distinct from all previously reported H3K4me1/me2/me3 recognition mechanisms^15, 16, 34^. The SDG8 CW domain binds to H3K4me1 with a pocket formed by one polar, two aromatic, and two hydrophobic residues^15^. The MORC3 CW domain binds to H3K4me1/me3 by two hydrophobic and two polar residues^34^. RDM15 specifically binds to H3K4me1 through the Tudor domain by an aromatic cage and a specific H-bond network^16^. The unshielded and exposed side of the H3K4me1 binding pockets in SDG8, MORC3, and RDM15 allows for the binding of H3K4me2 and H3K4me3, indicating a flexible mechanism for methylated H3K4 recognition. In contrast, the tandem PHD finger domain of Ehd3 forms a tight and narrow pocket only capable of accommodating H3K4me1, thereby restricting the binding of a greater number of methyl groups (Fig. 3). The PHD finger domain belongs to one of the most extensive families of domains present in reader proteins^35^, and recognizes multiple histone modifications ranging from H3K4 methylation and H3K14 acetylation^21^ to H3R2 methylation^36^. Most PHD finger domain structures are complexed with either an H3K4me2 or H3K4me3 peptide, exhibiting the conserved structural features that a single PHD finger domain forms an aromatic pocket for binding to a particular methylated lysine^22–25, 37–39^. However, the H3K4me1 binding pocket of Ehd3 is formed by the tandem PHD finger domain, which comprises two PHD fingers. A set of proteins indeed harbors a tandem PHD finger domain, but the recognition mechanisms for these proteins remain largely unclear. DPF3b and MOZ proteins recognize histones through their tandem PHD finger domains^21, 40–42^. In these proteins, the first PHD finger binds to H3K14ac^21, 40^ and H3K14ac/H3K14 crotonylation^41, 42^, respectively, whereas the second PHD finger interacts with unmodified H3K4, which represents a recognition mechanism distinct from Ehd3– H3K4me1.

In mammals, H3K4me1 marks enhancers but fewer H3K4me1 reader proteins have been identified to date^14^. The present analysis revealed the prevalence of Ehd3 homologs in plants and animals (Extended Data Fig. 8a). Distinct domain architecture and protein length were observed among Ehd3 homologs across plant and animal species (Extended Data Fig. 8a). Sequence alignment of the tandem PHD finger domains within Ehd3 homologs in land plants revealed a remarkable conservation of residues responsible for recognition of H3K4me1, particularly His441 and Try471 (Extended Data Fig. 8b). The ITC assays for the tandem PHD finger domains in Ehd3 homologs in *C. reinhardtii*, *P. patens*, *A. thaliana*, and *M. musculus* indicate their diverse binding affinities towards H3K4 methylations. Specifically, the tandem PHD finger domains in *M. musculus* showed binding affinity exclusively with the H3K4me0 peptide (PHD23, *K*_d_ = 6.5 μM; PHD56, *K*_d_ = 13.5 μM) (Extended Data Fig. 8c). The single PHD finger domain in *C. reinhardtii* failed to bind to any methylated H3K4. In contrast, the tandem PHD finger domain in *P. patens* exhibited a stronger affinity for H3K4me1, with even greater binding observed in *A. thaliana*, similar to that in *O. sativa* (Extended Data Fig. 8c). The tandem PHD finger domains within Ehd3 homologs in various species were predicted with AlphaFold (Extended Data Fig. 9)^43^; the interleaved topology and residues crucial for H3K4me1 binding were largely identical to those of Ehd3 in land plants, including *Zea mays*, *Triticum aestivum*, *A. thaliana*, and *Daucus carota* (Extended Data Fig. 9c–f). However, the tandem PHD finger domains in *P. patens* showed only partial similarity, and the overall conformations of those in *C. reinhardtii*, *penaeus vannamei*, *Elysia chlorotica*, *M. musculus*, and *H. sapiens* (Extended Data Fig. 9a–b, g–j) differed significantly from the tandem PHD finger domain within Ehd3. Collectively, the Ehd3 homolog in *C. reinhardtii*, lacking the tandem PHD finger domain structure, is unable to recognize any methylated H3K4. Upon transitioning from aquatic to terrestrial habitats, the Ehd3 homolog in the bryophyte *P. patens* evolved a tandem PHD finger domain capable of recognizing H3K4 methylation, especially H3K4me1. The tandem PHD finger domain within Ehd3 homologs in flowering plants exhibits binding affinity specific to H3K4me1 rather than H3K4me2. Thus, the tandem PHD finger domain initiates H3K4me1 recognition in land plants, and with highly exclusive affinity in flowering plants.

In this study, we observed that Ehd3 binds to H3K4me1 via its tandem PHD finger domain and interacts with the H3K36 methyltransferase SDG724 to deposit H3K36me2/me3 in the genome of rice. In Arabidopsis, the triple mutant *atx1/2/r7*, which lacked H3K4me1 deposition, exhibited a concomitant loss of both H3K4me1 and H3K36me3^44^. This observation supports the notion that H3K4me1 plays a critical role in directing H3K36me3 deposition in land plants. In addition, SDG725 is one of the H3K36 methyltransferases in rice and its CW domain recognizes methylated H3K4 with a preference for H3K4me1 over nonmethylated H3K4 or H3K4me2/me3^15^. We noted that SDG725 exhibited strong activity to catalyze H3K36 methylation when using H3K4me1 mononucleosomes as the substrate (Fig. 5g). Collectively, H3K4me1 readout is essential for H3K36 methyltransferases in land plants to deposit H3K36me2/me3, either through interactions with specific reader proteins or by employing their own H3K4me1-recognizing domain.

In yeast and animals, RNA polymerase II (RNAPII) recruits H3K36 methyltransferases to establish transcription elongation-coupled H3K36me3, and the Set2–Rpb1 interacting (SRI) domain within H3K36 methyltransferases is responsible for interaction with the C-terminal domain (CTD) of RNAPII^45, 46^. In contrast, we detected no SRI domains within the known H3K36 methyltransferases in higher plants (Extended Data Fig. 10a, b). Interestingly, the Arabidopsis H3K4me2-specific demethylase LDL3 possesses an SRI domain and interacts with the CTD-phosphorylated RNAPII^47^. Structure prediction by AlphaFold 3.0^48^ indicated that, similar to those within yeast Set2, the SRI domains within LDL3 homologs in land plants, including *P. patens*, *A. thaliana*, and *O. sativa*, display a conserved structure of a left-turned three-helix bundle (Extended Data Fig. 10c, d). We generated rice CRISPR/Cas9 mutants targeting *OsLDL3* in the NiP background. Two sgRNAs were designed to target the first exon of *OsLDL3*. One identified mutant, named *osldl3*, exhibited a 3-bp deletion at target site 1 and a 1-bp deletion at target site 2 (Extended Data Fig. 10e). Notably, deletion of Arabidopsis *AtLDL3*^47^ or rice *OsLDL3* leads to a decrease in H3K4me1 and an increase in H3K4me2 levels (Extended Data Fig. 10f, g). In addition, GST pull-down assays showed that Ser-2- or Ser-5-phosphorylated RNAPII was pulled down by the rice OsLDL3 SRI domain, but not by the H3K36 methyltransferases SDG724 or SDG725 (Extended Data Fig. 10h). The SRI domain of OsLDL3 especially interacted with the Ser-2-phosphorylated CTD of RNAPII, which marked transcription elongation, but not the nonphosphorylated CTD, indicating that RNAPII directs the transcription elongation-coupled recruitment of OsLDL3 (Extended Data Fig. 10i and Supplementary Tables 1–2). The foregoing observations suggest that, in plants, H3K4me1 deposition might be coupled with RNAPII, as RNAPII recruits H3K4me2 demethylase but not H3K36 methyltransferase. In yeast, H3K36me2/me3 suppress cryptic transcription initiation via Rpd3S-dependent histone deacetylation^49–51^, while in animals this suppression is mediated by Dnmt3b-dependent DNA methylation^52^. The Rpd3S complex asymmetrically engages with H3K36me3 nucleosomes through the chromodomains of the Eaf3-A/B subunits, positioning the catalytic center of Rpd3 towards the histone H4 N-terminal tail for deacetylation^53^. Dnmt3b recognizes H3K36me3 through the PWWP domain and binds to gene body regions to facilitate *de novo* DNA methylation^54^. Rpd3S-dependent histone deacetylation and Dnmt3b-dependent DNA methylation establish a repressive chromatin environment, which inhibits spurious entries of RNAPII and ultimately ensures the fidelity of transcription initiation. Plants and animals establish H3K36 methylation through different mechanisms; however, whether H3K36 methylation inhibits cryptic transcription initiation in plants remains to be elucidated.

As sessile organisms, plants must rapidly respond to environmental changes for survival, which is mainly achieved through dynamic regulation of gene transcription^55^. Thus, plants may undergo evolutionary adaptation to develop a more efficient mechanism to regulate the coordination among diverse histone modifications, thereby influencing the chromatin state and dynamically regulating gene transcription. Notably, H3K4 and H3K36 methylation are strongly associated with biotic and abiotic stresses in plants^56–62^. The present transcriptome data indicated that a set of stress-responsive genes were misregulated in the *ehd3-1*, *sdg724-1*, and *ehd3 sdg724* mutants (Extended Data Fig 5b, c). Thus, deposition of H3K36me2/me3 directed by H3K4me1 potentially plays an important role in plant environmental adaptation.

## Materials and Methods

### Plant material and growth conditions

The background rice material used in this study was *Oryza sativa* subsp. *japonica* ‘Nipponbare’ (NiP). Genomic DNA-edited mutants *ehd3-1*, *ehd3-4*, and *osldl3* were generated using the CRISPR/Cas9 system and obtained from BIORUN (Wuhan, China). The *sdg724-1* mutant was described previously^28^. The *ehd3 sdg724* double mutant was produced through a genetic cross between the single mutants *ehd3-1* and *sdg724-1*. The rice seedlings were cultured on Murashige and Skoog (MS) solid medium (comprising 2.2 g/L MS medium (M0222; Duchefa, Haarlem, The Netherlands), 3% (w/v) sucrose, and 0.4% (w/v) phytagel, adjusted to pH 5.8) under a 14-h photoperiod (30 °C light/28 °C dark) in an artificial climate chamber. Rice plants were grown in paddy fields at two distinct locations: Shanghai, under long-day (LD) conditions, and Sanya, under short-day (SD) conditions. For all experiments, at least 20 plants were used to evaluate the heading date, defined as the date at which the panicle emerged from the flag leaf. Arabidopsis seedlings were grown on MS solid medium (consisting of 4.9 g/L MS medium (M0255, Duchefa), 1% (w/v) sucrose, and 0.5% (w/v) agarose, adjusted to pH 5.7) under a 16-h photoperiod at 22 °C in an artificial climate chamber. *Chlamydomonas reinhardtii* strains were cultured in Tris-acetate-phosphate (TAP) medium^63^ on a rotary shaker at 25 °C in an artificial climate chamber. *Physcomitrella patens* were propagated on BCD medium^64^ under a 16-h photoperiod at 25 °C in an artificial climate chamber.

### Vector construction and transgenic plants

To generate the *Ubi::EGFP-Ehd3* construct, the *Ehd3* cDNA sequence was amplified by PCR and subsequently cloned into the pRHVnGFP vector^65^. For the *Ubi::YFP-SDG724* construct, the *SDG724* cDNA sequence was amplified by PCR and cloned into the pU1301 plant expression vector. For the *pEhd3::4×HA-Ehd3* and *pEhd3::4×HA-Ehd3mu* constructs, the native *Ehd3* promoter (2,005-bp upstream of ATG), the coding sequence for 4×HA, and the genomic DNA sequence of either wild-type *Ehd3* or the mutated *Ehd3* (Y471A and D496A) were amplified by PCR and cloned into the pCAMBIA1300 vector (CAMBIA, https://cambia.org). The primers used in cloning are listed in Supplementary Table 9.

The *Ubi::EGFP-Ehd3* and *Ubi::YFP-SDG724* constructs were transformed into NiP to generate transgenic overexpression plants, designated as *EGFP-Ehd3/NiP* and *YFP-SDG724/NiP*. In addition, *HA-Ehd3/ehd3-1* and *HA-Ehd3mu/ehd3-1* plants were generated by transforming the *pEhd3::4×HA-Ehd3* and *pEhd3::4×HA-Ehd3mu* constructs into the *ehd3-1* mutant, which lacks the transgene. Genetic transformations were conducted by BIORUN and stable transgenic plants were selected using hygromycin B (Roche, Mannheim, Germany). The stable transgenic plants were confirmed by western blotting using anti-GFP (ab290, Abcam, Cambridge, UK) or anti-HA (ab9110, Abcam) antibodies. In total, we obtained eight *EGFP-Ehd3/NiP* lines, five *YFP-SDG724/NiP* lines, two *HA-Ehd3/ehd3-1* lines, and two *HA-Ehd3mu/ehd3-1* lines. *EGFP-Ehd3/NiP-1* and *YFP-SDG724/NiP-1* were selected for the following experiments. In addition, *HA-Ehd3/ehd3-1-1*, *HA-Ehd3/ehd3-1-2*, and *HA-Ehd3mu/ehd3-1-1*, which demonstrated comparable abundance of HA-Ehd3 and HA-Ehd3mu proteins (Extended Data Fig. 4c), were selected for subsequent analysis.

### Phylogenetic analysis

The homologous proteins of Ehd3 were identified using the tandem PHD finger as bait in the UniProt database (https://www.uniprot.org). Sequence alignment and neighbor-joining dendrograms were constructed using MEGA version 10.2.6^66^.

### Protein expression and purification

The coding sequence of Ehd3 (1–563 amino acids (aa), 281–563 aa, 281–405 aa, 406– 563 aa, 406–473 aa, 473–563 aa, and 417–547 aa), OsLDL3 (1263–1832 aa), SDG724 (1–394 aa), and SDG725 (1240–1882 aa) were amplified by PCR using primers (Supplementary Table 9) and subsequently cloned into the pGEX-6P-1 vector (GE Healthcare, Milwaukee, WI, USA) to produce glutathione-*S*-transferase (GST)-fusion proteins. In addition, DNA fragments encoding Ehd3 (1–563 aa and 417–547 aa), the Ehd3 mutants (H441A, Y446A, Y471A, D486A, I489A, and D496A), OsLDL3 (1744– 1828 aa), SDG725 (1240–1898 aa), A0A2K3D188 (173–217 aa), A0A2K1L966 (364– 473 aa), Q1JPM3 (342–474 aa), Q8BRH4 (338–468 aa), and Q8BRH4 (911–1043 aa) were amplified by PCR using primers (Supplementary Table 9) and cloned into the pSUMO vector^67^. All recombinant proteins were expressed in *Escherichia coli* strain Rosetta (DE3) cells and induced with isopropyl-β-D-thiogalactoside. ZnCl_2_ was added during expression of the Ehd3, SDG724, and SDG725 proteins.

To purify GST-tagged proteins, cells were harvested and suspended in phosphate-buffered saline (1× PBS) containing 10% (v/v) glycerol, 0.1% (v/v) Triton X-100, 2 mM 1,4-dithiothreitol (DTT), and 0.1 mM ZnCl_2_, pH 7.4. After centrifuging the lysed cells, the supernatant was incubated with Glutathione Sepharose 4 Fast Flow (Cytiva, Uppsala, Sweden) at 4 °C for 2 h. Subsequently, the supernatants were discarded and the beads were washed three times with 1× PBS containing 10% (v/v) glycerol, 2 mM DTT, and 0.1 mM ZnCl_2_, pH 7.4. The GST-tagged proteins were eluted with an elution buffer (150 mM NaCl, 20 mM Tris-HCl, and 20 mM reduced glutathione, pH 7.4). The flow-through was concentrated and substituted with the buffer containing 150 mM NaCl, 20 mM Tris-HCl, 2 mM DTT, and 0.1 mM ZnCl_2_, pH 7.4, using an Amicon 10-KDa cutoff (Millipore, Burlington, MA, USA).

To purify the proteins cloned in the pSUMO vector, cells were harvested and suspended in the binding buffer (500 mM NaCl, 20 mM Tris-HCl, and 25 mM imidazole, pH 8.0). The lysed cells, processed using a high-pressure disruptor, were centrifuged, and the supernatant was loaded onto a Ni-NTA column (GE Healthcare). The proteins were gradually eluted using an elution buffer (500 mM NaCl, 20 mM Tris-HCl, and 500 mM imidazole, pH 8.0). Subsequently, the eluted proteins were incubated with ULP1 protease and then diluted for 3 h in buffer containing 500 mM NaCl and 20 mM Tris-HCl, pH 8.0. The diluted proteins were then loaded onto a Ni-NTA column, and the flow-through was further purified using a Superdex 75 column (GE Healthcare) equilibrated with buffer containing 100 mM NaCl, 20 mM Tris-HCl, and 1 mM Tris- (2-carboxyethyl)-phosphine hydrochloride (TCEP), pH 8.0. Finally, the proteins were concentrated using an Amicon 10-KDa cutoff (Millipore).

### Crystallization and structure determination

The purified Ehd3 protein (417–547 aa) was incubated with H3(1–12) K4me1 histone peptide (Supplementary Table 1) at a molar ratio of 1:1.2 at 4 °C for 1 h. Crystallization was performed by a Gryphon crystallization robot system with a sitting-drop vapor diffusion method. Specifically, 0.2 μl Ehd3 protein (10 mg/ml) and peptide mixture was combined with an equal volume of a crystallization kit solution (Hampton, Aliso Viejo, CA, USA) and incubated at 18 °C. The optimization procedure was performed using the hanging-drop vapor diffusion method at 18 °C. X-ray data for the crystals were collected at the B18U1 beamline of the Shanghai Synchrotron Radiation Facility and were processed using HKL3000^68^. The crystal structure was solved by the autosol and refined by the phenix.refine programs from Phenix^69^. The 2*F*_o_–*F*_c_ and *F*_o_–*F*_c_ electron density maps and unbiased omission map were regularly calculated and used as guides for the building of the H3K4me1 peptide, Zn^2+^ ions, and water molecules in COOT^70^. The data collection and refinement statistics are summarized in Supplementary Table 3. All molecular graphics were generated using PyMOL (https://pymol.org/).

### Structure prediction by AlphaFold

AlphaFold version 2.1.0^43^ was used to predict the structures of Ehd3 tandem PHD finger homologs, i.e., A0A2K3D188 (117–217 aa) in *Chlamydomonas reinhardtii*, A0A2K1L966 (364–464 aa) in *Physcomitrium patens*, A0A1D6LDK6 (401–498 aa) in *Zea mays*, A0A3B6RKR3 (432–532 aa) in *Triticum aestivum*, Q1JPM3 (354–450 aa) in *Arabidopsis thaliana*, A0A162A0W2 (424–519 aa) in *Daucus carota*, A0A3R7Q333 (793–889 aa) in *Penaeus vannamei*, A0A433TT05 (688–784 aa) in *Elysia chlorotica*, O14686 (1379–1475 aa) in *Homo sapiens*, and Q8BRH4 (914–1010 aa) in *Mus musculus*. AlphaFold version 3.0^48^ was used to predict the structures of SRI domain-containing homologs, i.e., A0A2K3DEA3 (1–1870 aa) in *Chlamydomonas reinhardtii*, A0A2K1LAV4 (1–1709 aa) and A0A2K1L6H4 (1–1814 aa) in *Physcomitrium patens*, SK05G17270.mRNA1 (1–1295 aa) in *Selaginella kraussiana*, Ceric.12G094400.1 (1– 1600 aa) in *Ceratopteris richardii*, F4JLS1 (1–1628 aa) in *Arabidopsis thaliana*, and Q336Y0 (1–1832 aa) in *Oryza sativa*. The protein codes correspond to the UniProt entries (https://www.uniprot.org), except for SK05G17270.mRNA1 and Ceric.12G094400.1, which were obtained from https://sk.ccgg.fun/blast.

### Peptide microarray

The MODified™ Histone Peptide Array (Active Motif, Carlsbad, CA, USA) was blocked with TTBS–milk buffer (20 mM Tris-HCl, 150 mM NaCl, 0.1% (v/v) Triton X-100, and 5% (w/v) non-fat dried milk, pH 7.4) at room temperature for 2 h. After washing three times with TTBS buffer (20 mM Tris-HCl, 150 mM NaCl, and 0.1% (v/v) Triton X-100, pH 7.4), the peptide array was incubated overnight at 4 °C with 500 nM GST-tagged full-length Ehd3 protein in TTBS–milk buffer. Subsequently, the peptide array was washed three times with TTBS buffer and incubated with the anti-GST antibody (M20007, Abmart, Shanghai, China) in the TTBS–milk buffer at room temperature for 1 h. After washing three times with TTBS buffer, the peptide array was incubated with the Goat Anti-Mouse IgG HRP antibody (M21001, Abmart) in the TTBS–milk buffer at room temperature for 1 h. The peptide array was washed three times with TBS buffer (20 mM Tris-HCl and 150 mM NaCl, pH 7.4) and imaged using a luminescence imaging workstation (Tanon, Shanghai, China). The results were analyzed using Array Analyze software (Active Motif).

### Peptide pull-down

Biotinylated histone peptides (2 μg) (H3(1–21), H3(1–21)K4me1, H3(1–21)K4me2, H3(1–21)K4me3, H3(1–21)K9me2, H3(1–21)K9ac, H3(1–21)K14ac, H3(21– 41)K27me3, H3(16–34)K27ac, and H3(21–41)K36me3) (Supplementary Table 1) were incubated with 20 μl prewashed streptavidin magnetic beads (NEB, Ipswich, MA, USA) at 4 °C for 2 h in 500 μl binding buffer (150 mM NaCl, 20 mM Tris-HCl, and 0.1% (v/v) Nonidet P-40). After washing three times, the peptide-bound beads were incubated with 2 μg GST-tagged full-length or truncated Ehd3 proteins at 4 °C for 2 h in 500 μl binding buffer. The beads were then washed three times with the binding buffer and boiled in 1× SDS loading buffer (50 mM Tris-HCl, 2% (w/v) sodium dodecyl sulfate, 0.01% (w/v) bromophenol blue, 2 mM DTT, and 2 mM β-mercaptoethanol, pH 6.8). The resulting products were subjected to SDS-PAGE and detected using the anti-GST antibody (M20007, Abmart).

### Semi-*in vivo* pull-down

Total nuclear proteins were extracted from wild-type NiP using a buffer containing 50 mM Tris-HCl, 5 mM MgCl_2_, 60 mM NaCl, 60 mM KCl, 2 mM DTT, and proteinase inhibitor cocktail (Roche), pH 7.5. The nuclear protein supernatant was incubated at 4 °C for 2 h with beads precoated with GST, GST-OsLDL3 SRI, GST-SDG724, or GST-SDG725 CW-SET proteins. After washing three times, the samples were analyzed by SDS-PAGE and western blotting using anti-Ser2P RNAPII (ab5095, Abcam) and anti-Ser5P RNAPII (ab5131, Abcam) antibodies.

### Isothermal titration calorimetry (ITC)

The ITC experiments were conducted using an iTC200 MicroCalorimeter (Malvern Panalytical, Malvern, UK) at the National Center for Protein Science Shanghai. The interactions involving the full-length Ehd3 (100 μM), wild-type or mutated Ehd3 truncations (417–547 aa, 100 μM), A0A2K3D188 (173–217 aa, 100 μM), A0A2K1L966 (364–473 aa, 100 μM), Q1JPM3 (342–474 aa, 100 μM), Q8BRH4 (338– 468 aa, 50 μM), and Q8BRH4 (911–1043 aa, 50 μM) were analyzed in the presence of the peptides H3(1–12)K4me0, H3(1–12)K4me1, H3(1–12)K4me2, or H3(1–12)K4me3 (1 mM), as well as OsLDL3 (1744–1828 aa, 100 μM) with the peptides NP CTD, S2P CTD, or S5P CTD (1 mM) (Supplementary Table 1). All reactions were performed at 25 °C in buffer containing 100 mM NaCl, 20 mM Tris-HCl, and 1 mM TCEP, pH 8.0. The titration data were fitted using Origin version 7.0 software (Supplementary Table 2).

### Immunoprecipitation followed by mass spectrometry (IP-MS)

Transgenic seedlings (2 g) were ground in liquid nitrogen and homogenized in lysis buffer (150 mM NaCl, 50 mM Tris-HCl, 5 mM MgCl_2_, 10% (v/v) glycerol, 0.1% (v/v) Nonidet P-40, 5 mM DTT, 0.1 mM PMSF, and proteinase inhibitor cocktail (Roche), pH 7.6). After sonication and centrifugation, the supernatant was incubated with the anti-GFP antibody (ab290, Abcam) at 4 °C for 3 h. The mixture was then incubated with protein A magnetic beads (Thermo Fisher Scientific, Waltham, MA, USA) at 4 °C for 3 h. Subsequently, the beads were washed with the same buffer and subjected to SDS-PAGE. The protein bands were then excised and subjected to in-gel digestion with trypsin. The resulting peptides were collected and analyzed using a nanoflow EASY-nLC 1200 system (Thermo Fisher Scientific) coupled with an Orbitrap Fusion Lumos mass spectrometer (Thermo Fisher Scientific). Raw data analysis was performed using Proteome Discoverer version 2.4 (Thermo Fisher Scientific) and an in-house Mascot server version 2.7 (Matrix Science, London, UK). The rice protein database was obtained from RGAP (http://rice.uga.edu). Data were searched using the following parameters: trypsin/P as the enzyme; allowing up to two missed cleavage sites; 10 ppm mass tolerance for MS and 0.05 Da for MS/MS fragment ions; carbamidomethylation on cysteine as fixed modifications; and protein N-terminal acetylation, oxidation on methionine, and N-terminal pyroQ as variable modifications. The incorporated Percolator in Proteome Discoverer was used for validation, accepting only the hits with false discovery rate ≤ 0.01 for the following analysis.

### GST pull-down

The GST or GST-SDG724 beads-coated protein was incubated with the full-length Ehd3 protein at 4 °C for 2 h in buffer containing 75 mM NaCl, 75 mM KCl, 50 mM HEPES, 5% (v/v) glycerol, 8 mM DTT, and 2 mM MgCl_2_, pH 8.0. After washing three times, the beads were boiled in 1× SDS loading buffer, subjected to SDS-PAGE, and stained with Coomassie Brilliant Blue.

### Yeast two-hybrid

The Ehd3 or SDG724 cDNA sequence was cloned into the pGADT7 and pGBKT7 vectors (Clontech, Kyoto, Japan), respectively, using specific primers (Supplementary Table 9). The resulting constructs were transformed into the Y2HGold yeast strain following the manufacturer’s instructions (WeidiBio, Shanghai, China). The interaction was screened on a synthetic defined medium lacking leucine, tryptophan, histidine, and adenine (SD/−L/−W/−H/−A).

### Co-immunoprecipitation (Co-IP)

Rice protoplast isolation and transformation were performed as described previously^71^. The *Ehd3* and *SDG724* cDNA sequences were cloned into the pRTVnMyc and pRTVnHA vectors^65^, respectively. After transformation of the constructs, the protoplasts were incubated at 28 °C for 20 h. The protoplasts were subsequently harvested in buffer containing 20 mM Tris-HCl, 150 mM NaCl, 1 mM EDTA, 2 mM DTT, 0.5% (v/v) Triton X-100, and proteinase inhibitor cocktail (Roche), pH 8.0. The harvested protoplasts were sonicated at 4 °C. After centrifugation, the supernatant was incubated with the agarose-conjugated anti-Myc antibody (M20012, Abmart) at 4 °C for 2 h. The immunoprecipitants were washed three times, subjected to SDS-PAGE, and detected with the anti-Myc (M20002, Abmart) and anti-HA (ab9110, Abcam) antibodies.

### Bimolecular fluorescence complementation (BiFC)

The *Ehd3* and *SDG724* cDNA sequences were cloned into the pRTVnVC and pRTVnVN vectors using the primers listed in Extended Data Table 4^65^, respectively. After transformation of the constructs, the protoplasts were incubated at 28 °C for 20 h. Fluorescence in the protoplasts was observed using an LSM 710 confocal laser scanning microscope (Carl Zeiss, Jena, Germany; https://www.zeiss.com).

### Histone methyltransferase assay

Either 500 nM recombinant Ehd3, GST-SDG724, GST-SDG724 mixed with Ehd3, or 100 nM SDG725 (CW-SET domain, 1240–1898 aa) were incubated with 1 μM unmodified (31467, Active Motif) or H3K4me1-modified (31585, Active Motif) mononucleosomes in histone methyltransferase buffer (50 mM Tris-HCl, 0.02% (v/v) Triton X-100, 1 mM TCEP, and 160 μM SAM, pH 8.5) at room temperature for 6–24 h. Reactions were stopped by adding 1× SDS loading buffer, then were subjected to SDS-PAGE and detected by western blotting using the anti-H3K36me2 (ab9049, Abcam) and anti-H3K36me3 (ab9050, Abcam) antibodies.

### RNA extraction, library construction and sequencing

Fourteen-day-old *Oryza sativa* and *Arabidopsis thaliana* seedlings (100 mg) grown on MS medium, 7-day-old *Physcomitrella patens* grown on BCD medium, and *Chlamydomonas reinhardtii* cultured in TAP medium to a density of 4–8 × 10^6^ cells/ml were used for mRNA extraction. The mRNA was extracted using the RNAprep Pure Plant Kit (Tiangen Biotech, Beijing, China). Three independent biological replicates of RNA-seq libraries were prepared using the KAPA Stranded mRNA-seq Kit (Kapa Biosystems, Wilmington, MA, USA) and sequenced on an Illumina NovaSeq 6000 instrument by NEO BIO (Shanghai, China). Raw reads were cleaned with Cutadapt version 3.5^72^ to remove low-quality bases and sequencing adapters. Trimmed reads were aligned to the reference genomes of different species using HISAT2 version 2.1.0^73^. Samtools version 1.17^74^ was used to remove reads with low mapping quality (MAPQ < 20), and then featureCounts version 2.0.1^75^ was used to quantify the number of reads mapped to exons. For rice RNA-seq, the DESeq2 version 1.36.0^76^ package for R (version 4.2.1) was used to screen DEGs (|fold of change| ≥ 1.5 and adjusted *p*-value ≤ 0.05). Gene ontology (GO) functional enrichment analysis for specific gene sets was conducted through the online platform CARMO^77^.

The reference genomes used for rice and Arabidopsis were MSU7 (http://rice.uga.edu/) and TAIR10 (https://www.arabidopsis.org/), respectively. The reference genomes for other species were obtained from Ensembl (https://www.ensembl.org/). The genomes correspond to specific species: R64-1-1 for *Saccharomyces cerevisiae*, BDGP6.32 for *Drosophila melanogaster*, GRCm39 for *Mus musculus*, GRCh38.p14 for *Homo sapiens*, Chlamydomonas_reinhardtii_v5.5 for *Chlamydomonas reinhardtii*, and Phypa_V3 for *Physcomitrella patens*.

### Chromatin immunoprecipitation (ChIP)

Fourteen-day-old *Oryza sativa* and *Arabidopsis thaliana* seedlings (2 g) grown on MS medium, and 7-day-old *Physcomitrella patens* grown on BCD medium, were fixed in cross-linking buffer (0.4 M sucrose, 10 mM Tris-HCl, 1 mM PMSF, 1 mM EDTA, and 1% (v/v) formaldehyde, pH 8.0). *Chlamydomonas reinhardtii* grown in 1 L TAP medium to a density of 4–8 × 10^6^ cells/ml were harvested and fixed in cross-linking buffer (20 mM HEPES-KOH, 80 mM KCl, and 0.4% (v/v) formaldehyde, pH 7.6). The ground seedlings or cells were resuspended in isolation buffer (0.25 M sucrose, 15 mM PIPES, 5 mM MgCl_2_, 60 mM KCl, 15 mM NaCl, 1 mM CaCl_2_, 0.9% (v/v) Triton X-100, 1 mM PMSF, and proteinase inhibitor cocktail (Roche), pH 6.8). Chromatin was sonicated in lysis buffer (50 mM HEPES, 150 mM NaCl, 1 mM EDTA, 1% (w/v) SDS, 0.1% (w/v) sodium deoxycholate, 1% (v/v) Triton X-100, and proteinase inhibitor cocktail (Roche), pH 7.5) using a Bioruptor™ Plus sonication device (Diagenode, Liege, Belgium). After centrifugation, the supernatant was incubated with antibodies including anti-H3K4me1 (ab8895, Abcam), anti-H3K4me2 (07-030, Millipore), anti-H3K4me3 (07-473, Millipore), anti-H3K36me1 (ab9048, Abcam), anti-H3K36me2 (ab9049, Abcam), anti-H3K36me3 (ab9050, Abcam), anti-H3 (ab1791, Abcam), anti-HA (ab9110, Abcam), or anti-GFP (ab290, Abcam) overnight at 4 °C. The prewashed protein A magnetic beads (Thermo Fisher Scientific) were added to the mixture and incubated at 4 °C for 2 h. The immunoprecipitates were washed sequentially: once with low-salt buffer (150 mM NaCl, 20 mM Tris-HCl, 0.1% (w/v) SDS, 1% (v/v) Triton X-100, and 2 mM EDTA, pH 8.0), once with high-salt buffer (500 mM NaCl, 20 mM Tris-HCl, 0.1% (w/v) SDS, 1% (v/v) Triton X-100, and 2 mM EDTA, pH 8.0), once with LiCl buffer (10 mM Tris-HCl, 0.25 M LiCl, 1% (w/v) sodium deoxycholate, 1% (v/v) Nonidet P-40, and 1 mM EDTA, pH 8.0), and twice with Tris-EDTA buffer (10 mM Tris-HCl and 1 mM EDTA, pH 8.0). The immunoprecipitates were eluted with elution buffer (0.5% (w/v) SDS and 0.1 M NaHCO_3_), followed by reverse crosslinking overnight at 65 °C. Subsequently, the samples were treated with RNase A and proteinase K. DNA extraction and ChIP-seq library construction were conducted as described previously^78^.

### ChIP sequencing (ChIP-seq) analysis

DNA fragments for ChIP assays were prepared for sequencing on an Illumina NovaSeq 6000 instrument by NEO BIO (Shanghai, China). Raw reads trimmed by Cutadapt were aligned to genomes using Bowtie2 version 2.4.5^79^. Reads with low-quality mapping (MAPQ < 20) and duplicate reads were filtered out using Samtools version 1.17^74^. Macs3 version 3.0.0b3^80^ and MAnorm v1.3.0^81^ were used for peak-calling and differential peaks detection, respectively. The ChIPpeakAnno version 3.30.1^82^ package for R version 4.2.1 was used to annotate peaks to genes. BEDtools version 2.31.0^83^ and deepTools version 3.5.3^84^ were used to calculate the normalized signals of fragments aligned to genes or peaks. Genomic track files generated by deepTools version 3.5.3^84^ were visualized in the genome browser IGV version 2.11.1^85^. Metagene plots and heatmaps for regions from −3 kb to 3 kb around genes were plotted using the computeMatrix and plotHeatmap tools in deepTools version 3.5.3^84^. Each ChIP-seq sample included at least two independent biological replicates.

### Reporting summary

Further information on research design is available in the Nature Portfolio Reporting Summary linked to this article.

### Data and code availability

The data supporting the findings of this work are available within the main text, supplementary materials, the listed Protein Data Bank (PDB) accession, or the listed NCBI GEO accessions. The atomic coordinates of the Ehd3-bound H3K4me1 peptide structure have been deposited in the PDB (accession code 8xr1). The RNA-seq and ChIP-seq (anti-H3K4me1/me2/me3, anti-H3K36me1/me2/me3, anti-H3, anti-HA, and anti-GFP) data for *C. reinhardtii*, *P. patens*, and *O. sativa* in this work have been deposited in the National Center for Biotechnology Information (NCBI) database (accession number GSE266911). The ChIP-seq data for *O. sativa* used in correlation analyses were obtained from the NCBI database (accession numbers GSE79033, GSE126436, and GSE109616). The *H. sapiens* ChIP-seq data used in correlation analyses were obtained from the NCBI database (accession numbers GSE175752, GSE179461, GSE107599, and GSE200770). The RNA-seq and ChIP-seq data for H3K4 and H3K36 methylations were obtained from the NCBI database (accession numbers GSE73407 and GSE242874 for *S. cerevisiae*; GSE47281, GSE55555, GSE47285, GSE118785, and SRR1197331 for *D. melanogaster*; GSE95781 and GSE118785 for *M. musculus*; GSE51176, GSE169207, and GSE174252 for *H. sapiens*; and GSE258772 and GSE188493 for *A. thaliana*). The RNA-seq and ChIP-seq data for H3K4 methylations in the *A. thaliana atldl3* mutant were obtained from the DRA (accession number DRA008014) and NCBI (accession number PRJNA934737) databases. Related code is available from the lead contact on reasonable request.

## Acknowledgements

This work was supported by the National Key Research and Development Program of China (2024YFE0105100), the National Natural Science Foundation of China (NSFC31930017, NSFC31800207, NSFC32100453, and NSFC32400467), and the Open Research Fund of State Key Laboratory of Crop Gene Exploration and Utilization in Southwest China, Sichuan Agricultural University (SKL-KF202412). We thank the Shanghai Synchrotron Radiation Facility for providing technical support and assistance with crystallization data collection; the National Center for Protein Science Shanghai for assistance in ITC experiments; and Robert McKenzie, PhD, from Liwen Bianji (Edanz) (www.liwenbianji.cn) for editing a draft of this manuscript.

## Author contributions

A.D. conceived and designed the research. A.D., B.L., and J.G. supervised the experiments and interpreted the data; J.Wu, K.D., Y.L., C.H., Q.X., X.Z., Z.W., F.L., and J.G. collected and analyzed the data; J.Wang, W.X., X.L., and X.W. analyzed the bioinformatic data; J.Wu, J.Wang, K.D., Y.L., W.-H.S., J.G., B.L., and A.D. wrote the manuscript.

## Competing interests

The authors declare no competing interests.

## Notes

### Competing Interest Statement

The authors have declared no competing interest.

