## Supplemental Figures for "H3K4me1 directs H3K36me2 and H3K36me3 deposition in land plants"

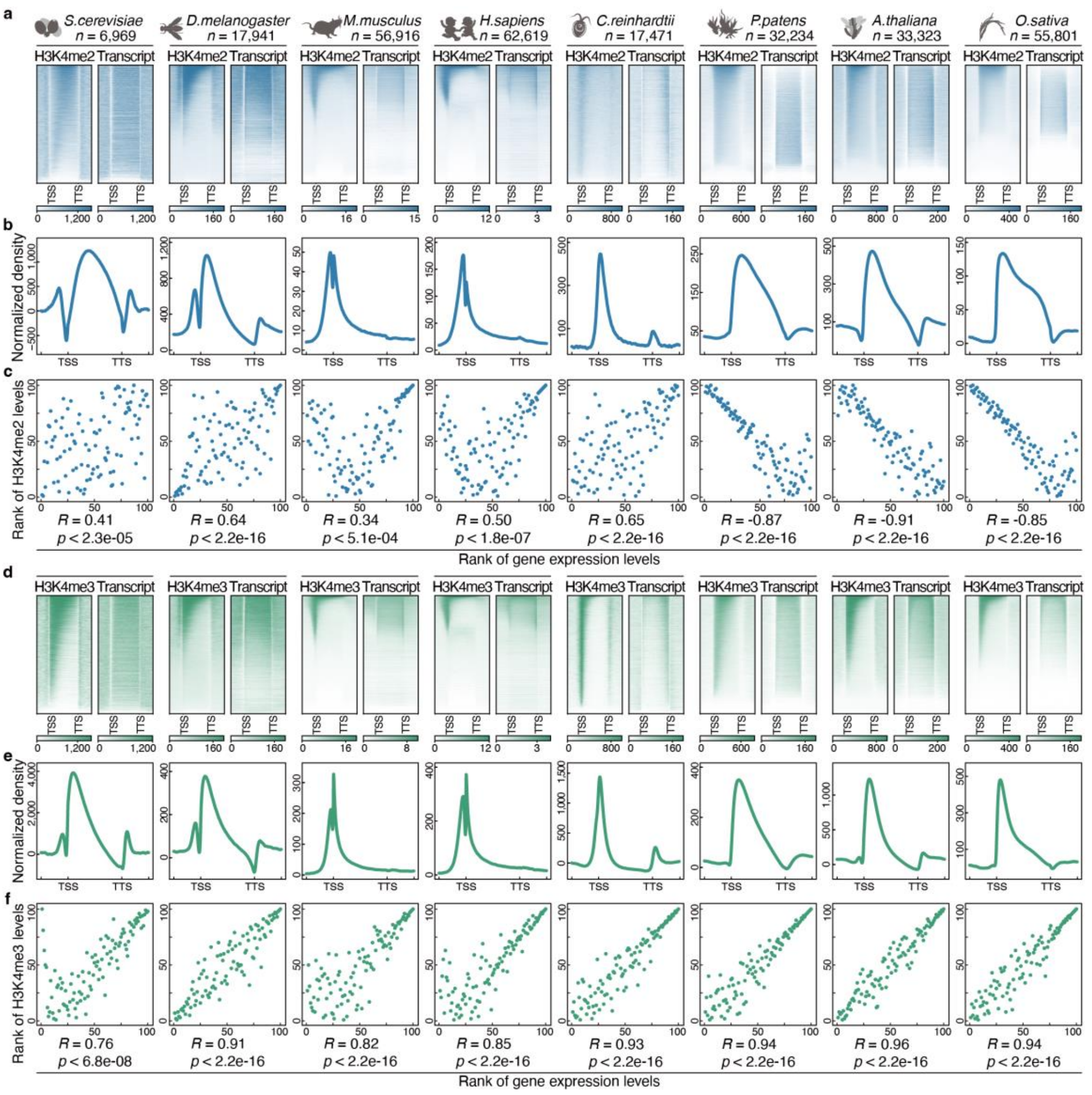


**Extended Data Fig. 1: Characterization of the genome-wide distributions of H3K4me2 and H3K4me3 across diverse species.**

**a**,**d**, Heatmaps showing the individual gene distribution of H3K4me2 (a) and H3K4me3 (d) (reads per kilobase per million mapped reads; RPKM), together with their associated transcription level (RPKM) in eight species. All genes are sorted based on the enrichment of H3K4me2 (a) or H3K4me3 (d). **b**,**e**, Integrative genomic distributions of H3K4me2 (b) and H3K4me3 (e) in various species. **c**,**f**, Scatter plots presenting the correlations of H3K4me2 (c) and H3K4me3 (f) levels with gene transcription levels in eight species. The species from left to right are *Saccharomyces cerevisiae*, *Drosophila melanogaster*, *Mus musculus*, *Homo sapiens*, *Chlamydomonas reinhardtii*, *Physcomitrium patens*, *Arabidopsis thaliana*, and *Oryza sativa*, respectively. The transcribed genes enriched with H3K4me2 or H3K4me3 modifications were divided into 100 groups based on the levels of their transcription and modification. The average values of each group of modifications and transcription were calculated for analysis. The heatmaps and plots present the region from 3 kb upstream of the transcription start site (TSS) to 3 kb downstream of the transcription termination site (TTS). For scatter plots, Spearman’s rank correlation coefficient indicates the correlation between the methylation level and gene transcription level. The *p*-value was determined based on Spearman’s rank correlation test.

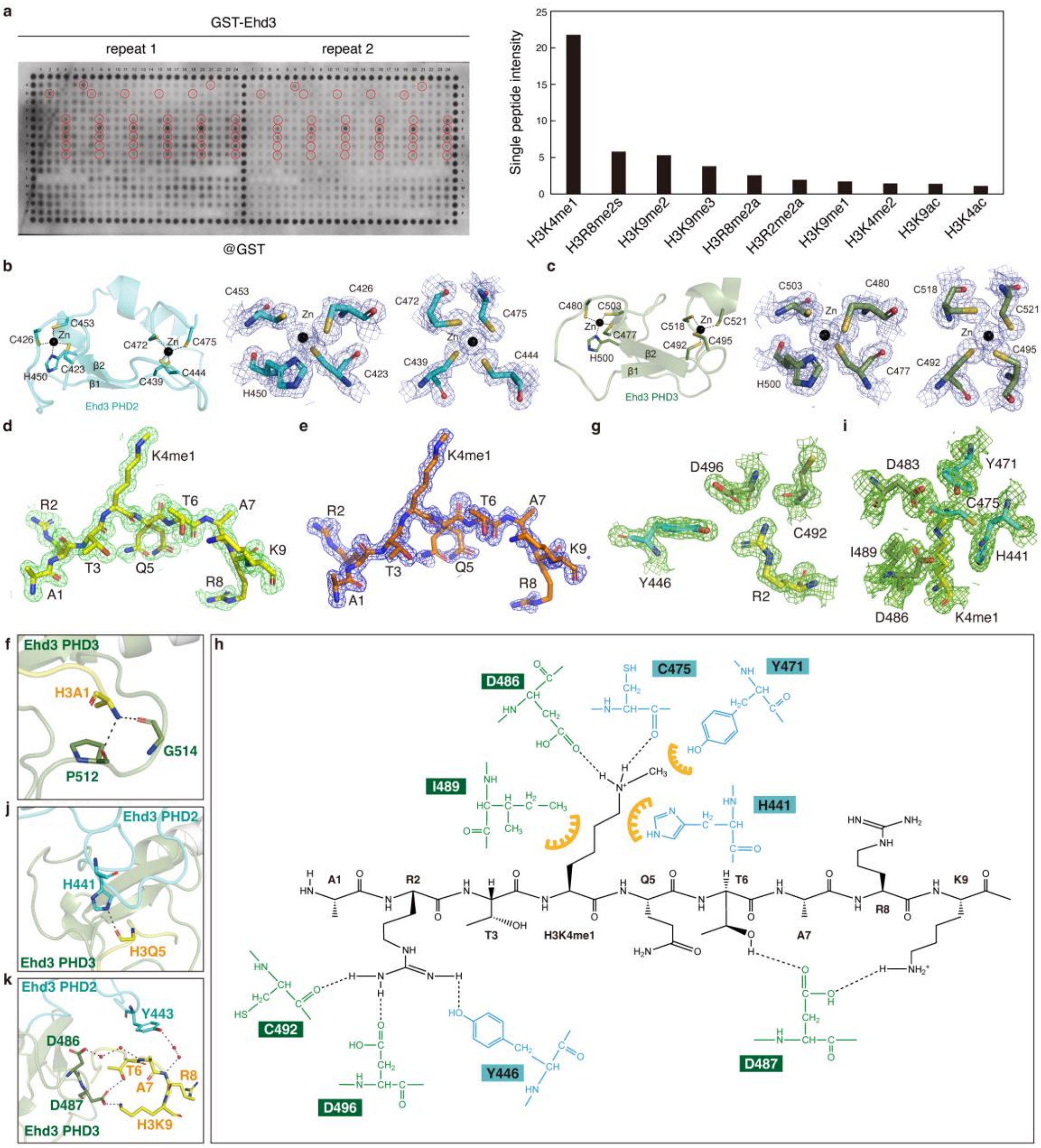


**Extended Data Fig. 2: Ehd3 recognizes H3K4me1 peptide.**

**a**, Representative array images showing histone binding preferences of glutathione-*S*-transferase (GST)–Ehd3 on a histone peptide microarray. Red circles indicate the locations of H3K4me1-modified peptides. Histogram showing the relative binding intensity of GST–Ehd3 to the indicated modified histone peptide. **b**,**c**, Critical residues of Ehd3 plant homeodomain 2 (PHD2) (b) and PHD3 finger (c) involved in zinc ion binding are shown as sticks. Zinc ions are depicted as spheres in black. The 2$F_o$–$F_c$ electron density maps are contoured at 1.5 sigma levels. **d**, Overall conformation of the bound H4K3me1 peptide. The 2$F_o$–$F_c$ electron density map of the peptide is contoured at 1.0 sigma levels. **e**, Unbiased omission map of the bound H4K3me1 peptide. The $2F_o–F_c$ electron density map of the peptide is contoured at 3.0 sigma levels. **f**, Detailed interaction of Ehd3 with H3A1. **g**, Residues responsible for H3R2 recognition. The $2F_o–F_c$ electron density maps of the peptide are contoured at 1.5 sigma levels. **h**, Schematic representation showing the intermolecular interactions between Ehd3 and the side chains of H3K4me1 peptide residues. The hydrogen bonds are highlighted by dashed lines. **i**, Residues responsible for H3K4me1 recognition. The $2F_o–F_c$ electron density maps of the peptide are contoured at 1.5 sigma levels. **j**,**k**, Detailed interaction of Ehd3 with H3Q5 (j) and with H3T6, H3A7, H3R8, and H3K9 (k).

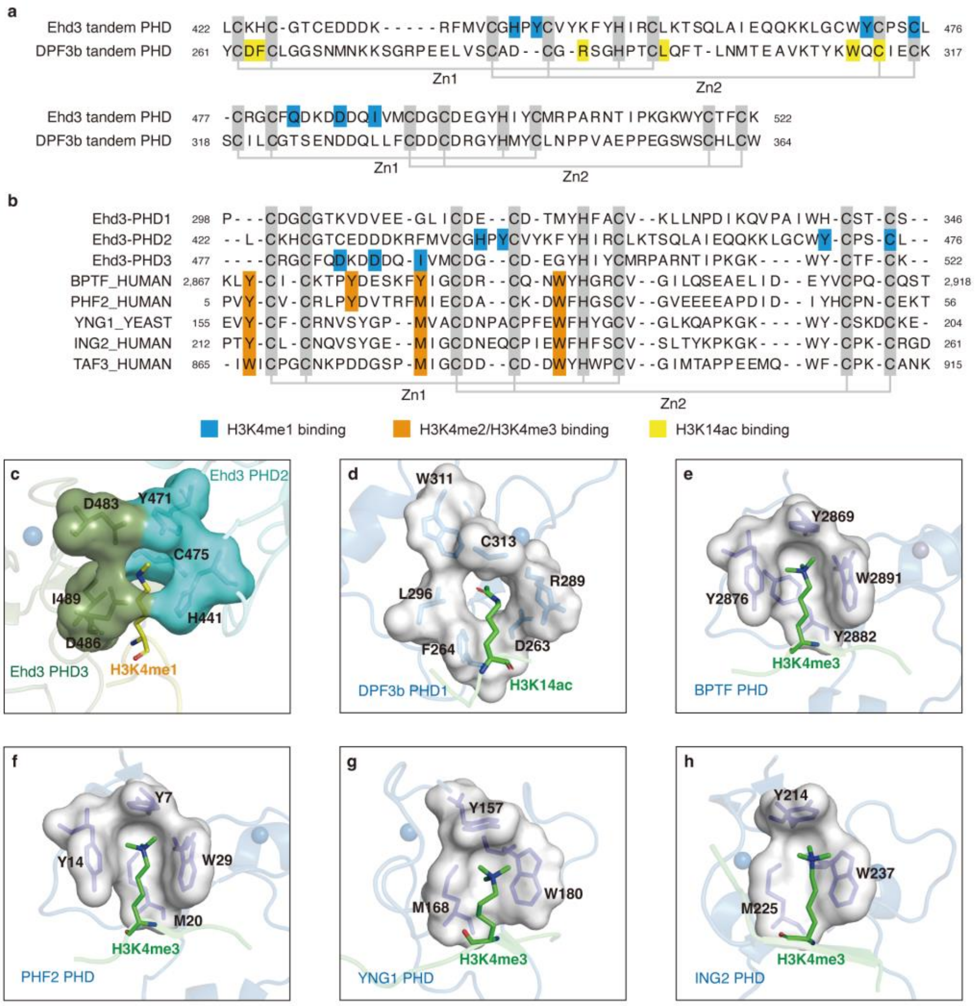


**Extended Data Fig. 3: Structure-based characterization of the representative PHD finger.**

**a**,**b**, Sequence alignment of the tandem plant homeodomain (PHD) of Ehd3 with the tandem PHD of DPF3b (a), and PHD fingers in Ehd3 with other PHD-containing proteins (b). The critical residues of PHD fingers are classified into three groups, distinguished by different colors as indicated. Zinc-coordinating residues are shaded in light grey. **c**–**h**, Representative structure of the PHD domain including Ehd3 (c), DPF3b (d), BPTF (e), PHF2 (f), YNG1 (g), and ING2 (h), in complex with various H3 modifications. The PDB structures include: 5szb (DPF3b), 2f6j (BPTF), 3kqi (PHF2), 2jmj (YNG1), and 2g6q (ING2).

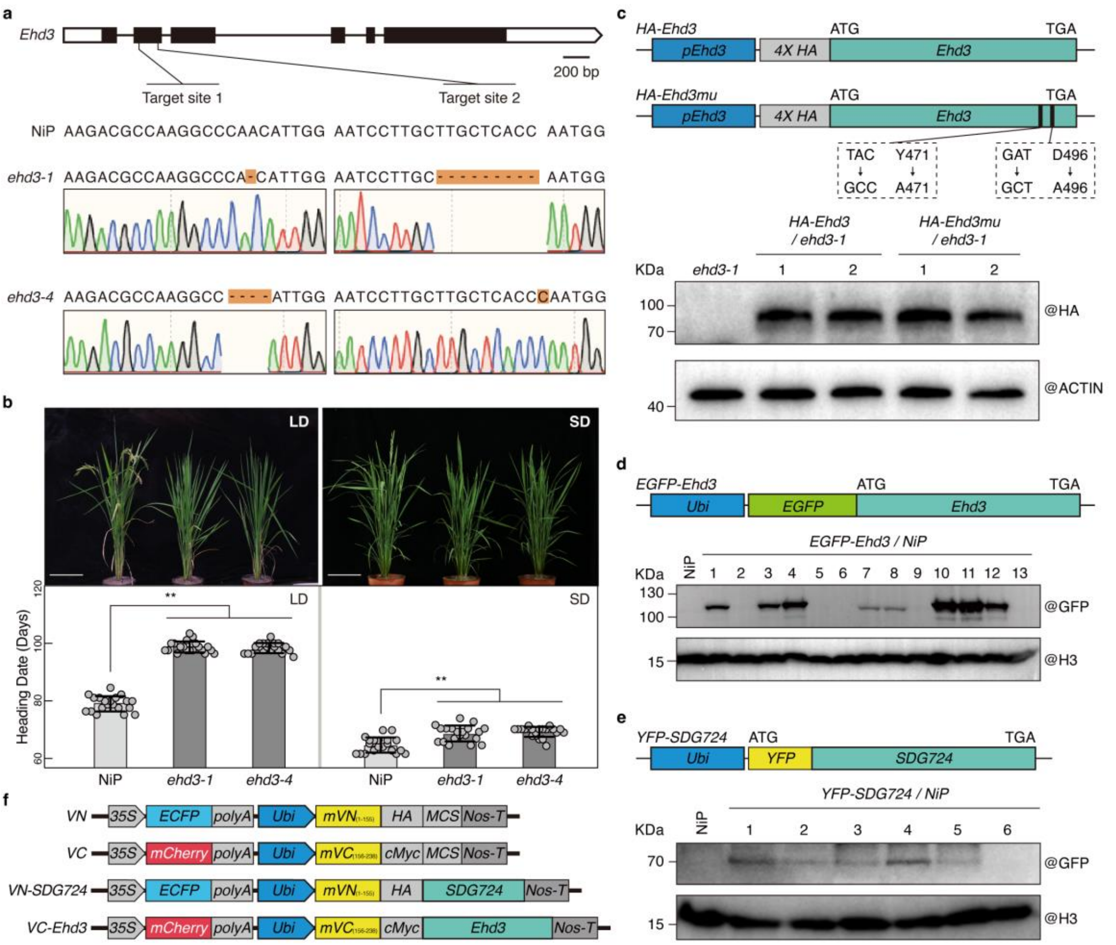


**Extended Data Fig. 4: Generation of CRISPR/Cas9 mutants and transgenic plants involving *Ehd3* and *SDG724*.**

**a**, Identification of the *ehd3* mutants generated by CRISPR/Cas9 editing. The untranslated regions, exons, and introns of *Ehd3* are indicated by blank rectangles, black rectangles, and black lines, respectively. The altered sequences in the *ehd3-1* and *ehd3-4* mutants are highlighted in orange. **b**, Representative images showing wild-type rice 'Nipponbare' (NiP), *ehd3-1*, and *ehd3-4* plants grown under long-day (LD) and short-day (SD) conditions, respectively. Scale bar, 20 cm. Histograms show days to heading in NiP, *ehd3-1*, and *ehd3-4* plants under LD and SD conditions. Values are shown as means ± standard deviation of 20 individual plants. Statistical significance was determined by a one-way ANOVA test: ** *p*-value < 0.01. **c**, Schematic representation showing the *pEhd3::4×HA-Ehd3* (*HA-Ehd3*) and *pEhd3::4×HA-Ehd3mu* (*HA-Ehd3mu*) constructs. The western blot shows comparable levels of HA-Ehd3 and HA-Ehd3mu protein in the transgenic lines. **d**,**e**, Schematic representation showing the *EGFP-Ehd3* (D) and *YFP-SDG724* (E) constructs. The western blot shows the EGFP-Ehd3 (D) and YFP-SDG724 (E) protein in the different transgenic lines. **f**, Schematic representation of vectors for bimolecular fluorescence complementation assays. The mVN (1–155 amino acids (aa)) and mVC (156–238 aa) are the N- and C-terminals of mVenus, respectively. To evaluate plasmid transfection and gene expression efficiency, the marker genes *ECFP* and *mCherry* are incorporated into the vectors.

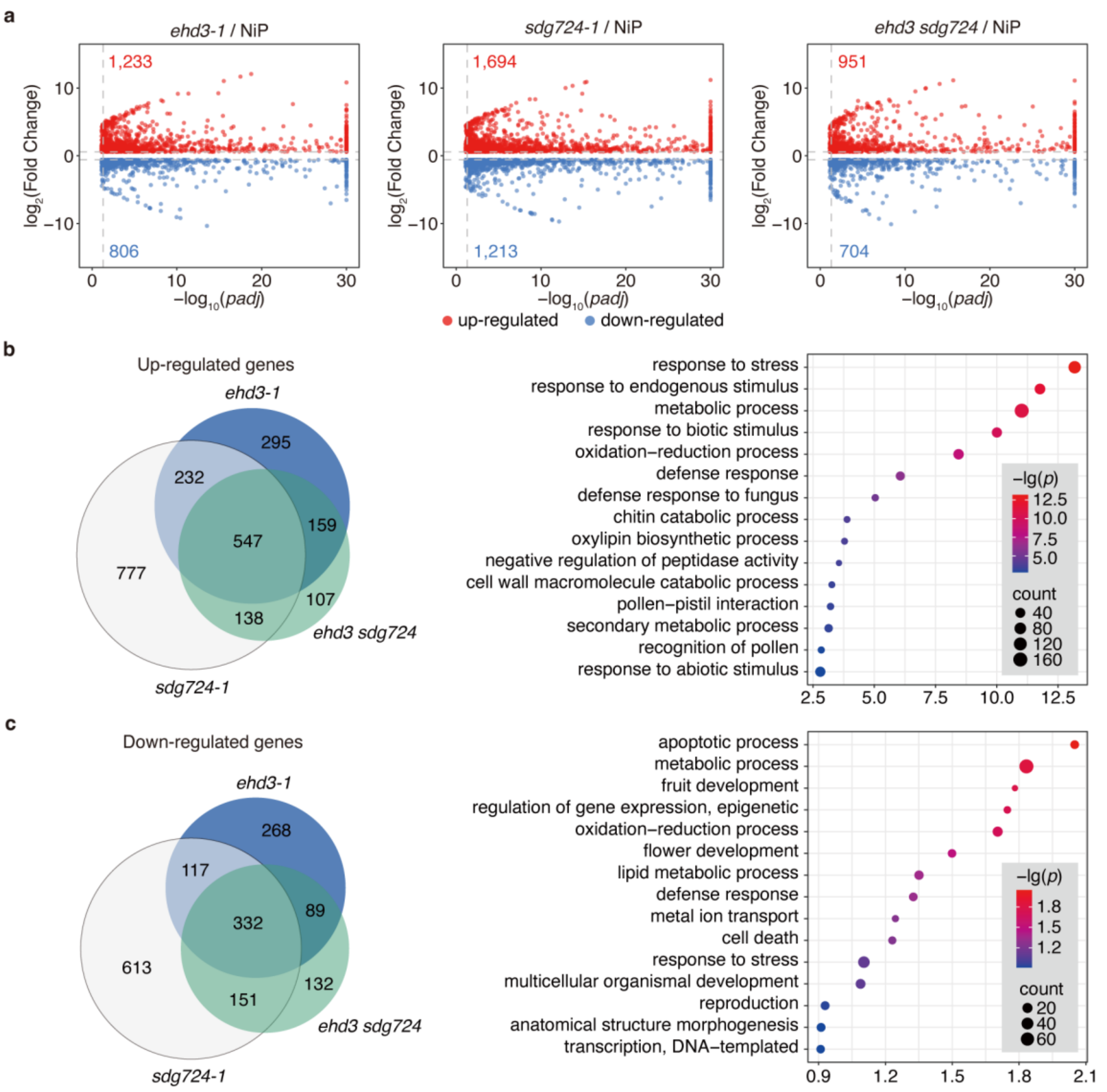


**Extended Data Fig. 5: Ehd3 and SDG724 coregulate expression of a set of genes.** **a**, Volcano plots showing differentially expressed genes identified by RNA-seq analysis in the *ehd3-1*, *sdg724-1*, and *ehd3 sdg724* mutants compared with the wild-type rice ‘Nipponbare’ (NiP). The *x*-axis indicates the adjusted *p*-value, and the *y*-axis indicates the fold change in transcription. Up- and down-regulated genes (red and blue dots, respectively) were identified based on changes and adjusted *p*-value above thresholds (above noise and $\geqslant$ 1.5-fold change, shown as dashed lines) and with the *p*-value $\leqslant$ 0.05. The adjusted *p*-value was calculated using Benjamini–Hochberg correction. **b**,**c**, Venn diagrams showing the overlap of up-regulated (b, left) or down-regulated (c, left) genes in the *ehd3-1*, *sdg724-1*, and *ehd3sdg724* mutant plants. Dotplots enrichment map showing biological processes associated with up-regulated genes (b, right) or down-regulated genes (c, right). The top 15 catalogs by *p*-value are presented. The color of the dot depends on the *p*-value, and the size is determined by the respective gene number.

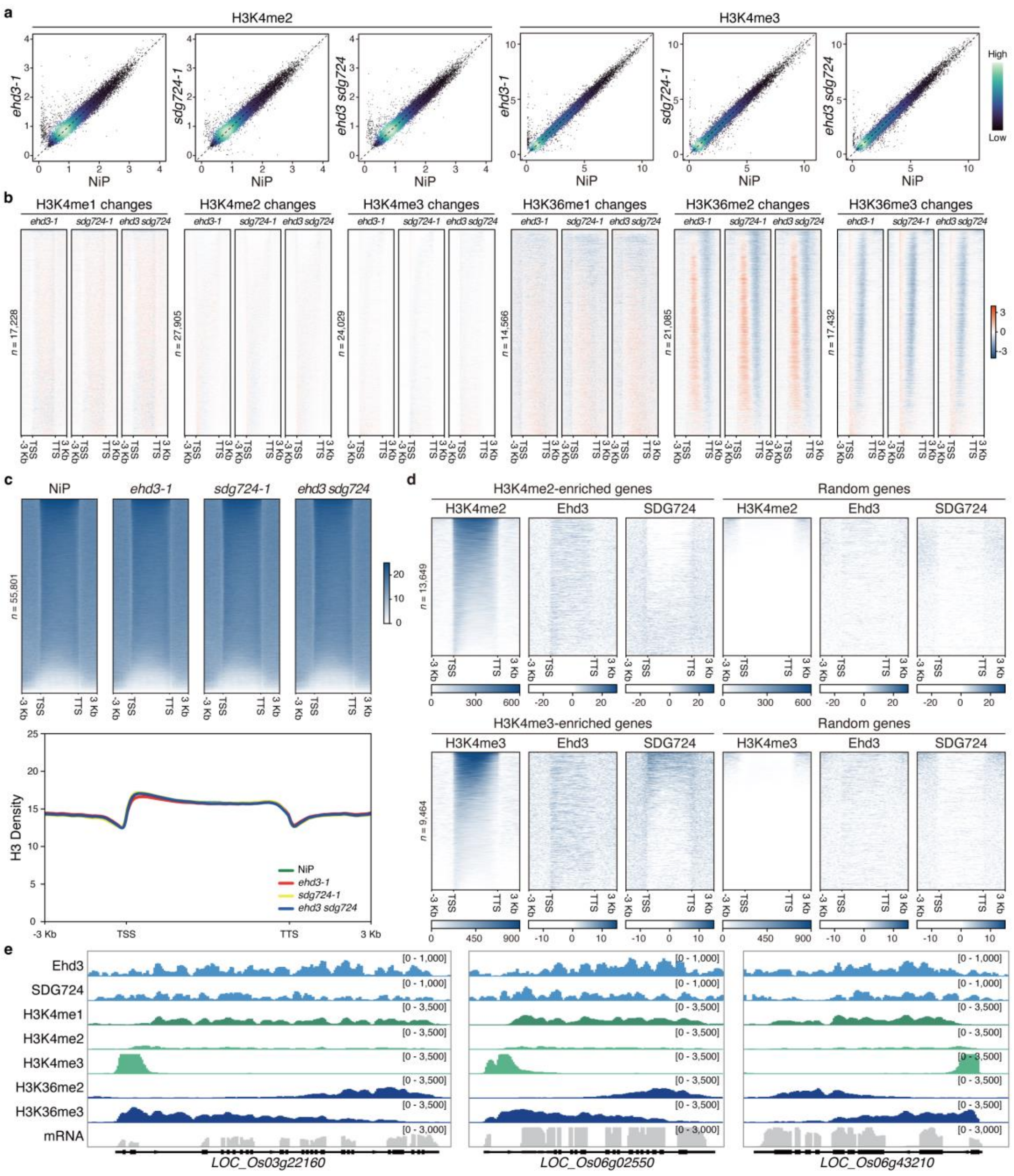


**Extended Data Fig. 6: Genome-wide changes of H3K4/H3K36 methylations and H3 levels in the mutants, as well as the Ehd3/SDG724 levels within H3K4me2- and H3K4me3-enriched genes.**

**a**, Scatter plots comparing H3K4me2 and H3K4me3 between the wild-type rice 'Nipponbare' (NiP), *ehd3-1*, *sdg724-1*, and *ehd3 sdg724* mutants. Each dot represents the histone methylation level of one peak corrected with MAnorm. The transition from deep blue to cyan indicates the increase of dot density. **b**, Heatmaps showing the changes of H3K4me1, H3K4me2, H3K4me3, H3K36me1, H3K36me2, and

H3K36me3 levels (bins per million mapped reads; BPM) in the *ehd3-1*, *sdg724-1*, and *ehd3 sdg724* mutants, respectively, compared with NiP, from 3 kb upstream of the transcription start site (TSS) to 3 kb downstream of the transcription termination site (TTS). All genes are sorted based on the enrichment of the corresponding histone methylation identified in NiP. Numbers of genes are shown on the left. **c**, Heatmaps and metaplots showing the H3 distribution (reads per kilobase per million mapped reads; RPKM) from 3 kb upstream of the TSS to 3 kb downstream of the TTS in NiP, *ehd3-1*, *sdg724-1*, and *ehd3 sdg724*. All genes in heatmaps are sorted based on enrichment of H3. Numbers of genes are shown on the left. **d**, Heatmaps showing the enrichment (RPKM) of Ehd3 or SDG724 within H3K4me2 without H3K4me1-enriched genes and H3K4me3 without H3K4me1-enriched genes from 3 kb upstream of the TSS to 3 kb downstream of the TTS. An equal number of genes lacking H3K4me1/H3K4me2 and H3K4me1/H3K4me3 enrichment were randomly selected as a control. All genes are sorted based on the enrichment of H3K4me2 or H3K4me3 identified in NiP. Numbers of genes are shown on the left. **e**, Integrative Genomics Viewer images of ChIP-seq and RNA-seq data showing the distributions of Ehd3, SDG724, H3K4me1, H3K4me2, H3K4me3, H3K36me2, H3K36me3, and mRNA along the representative genes in NiP.

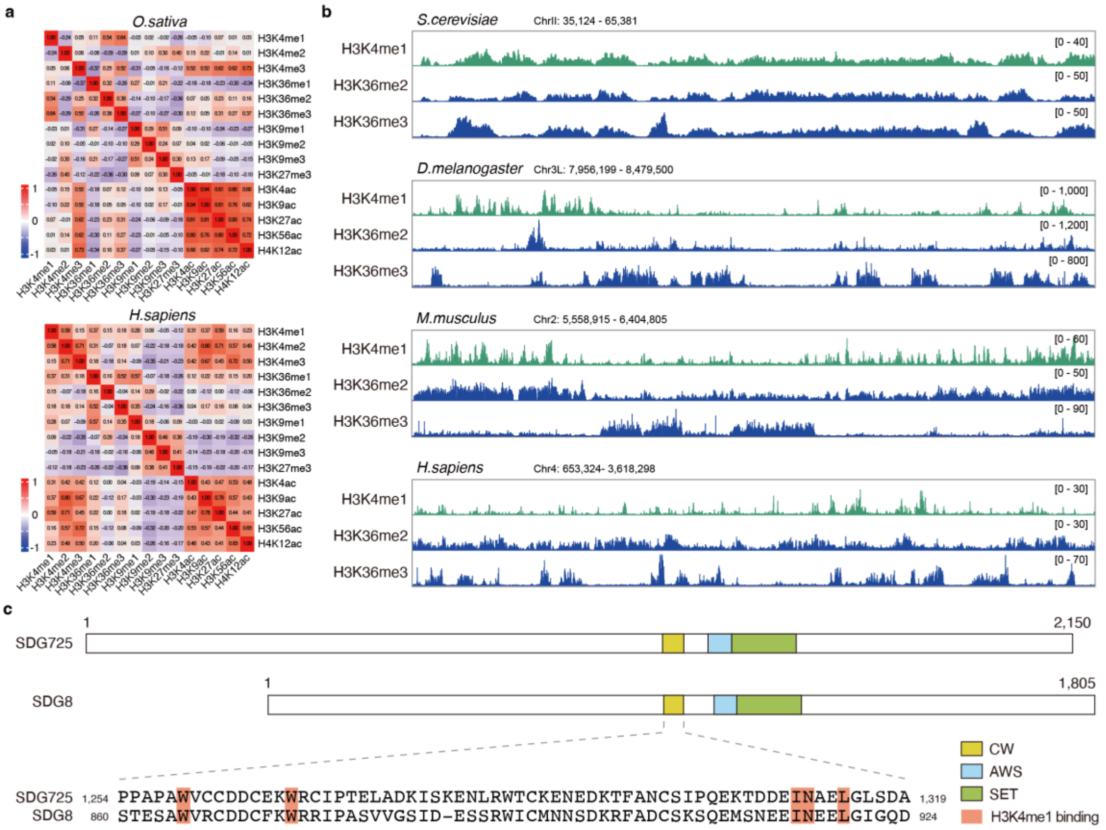


**Extended Data Fig. 7: H3K4me1 directs H3K36me2 and H3K36me3 deposition in plants.**

**a**, Spearman correlation heatmap of histone modifications in *Oryza sativa* and *Homo sapiens*. **b,** Integrative Genomics Viewer images of ChIP-seq data showing the H3K4me1, H3K36me2, and H3K36me3 profile plots of the representative region in *Saccharomyces cerevisiae*, *Drosophila melanogaster*, *Mus musculus*, and *Homo sapiens***. c,** Schematic representation showing similar protein domain architecture of rice SDG725 and Arabidopsis SDG8. Sequence alignment of the CW domain showing the conserved residues responsible for H3K4me1 binding, highlighted in orange rectangles.

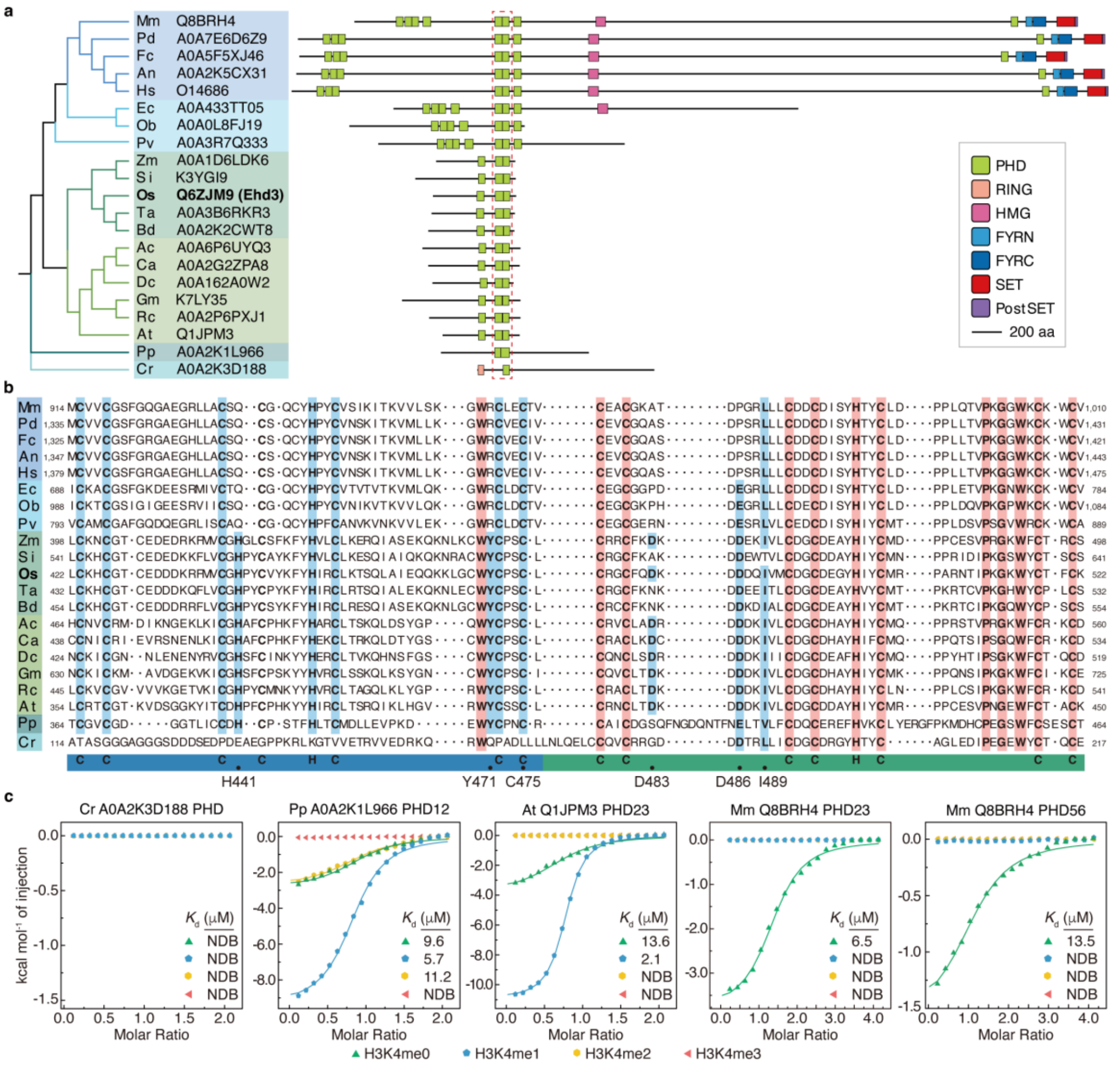


**Extended Data Fig. 8: Ehd3 homologs selectively recognize H3K4me1 in land plants.**

**a**, Neighbor-joining dendrogram (left) and protein domain architecture (right) of the tandem plant homeodomain (PHD) domain-containing proteins in higher eukaryotes. The dendrogram was constructed based on the Ehd3 tandem PHD sequence alignment. The diverse protein domains are displayed in different colored boxes as indicated. Mm, *Mus musculus*; Pd, *Phyllostomus discolor*; Fc, *Felis catus*; An, *Aotus nancymaae*; Hs, *Homo sapiens*; Ec, *Elysia chlorotica*; Ob, *Octopus bimaculoides*; Pv, *Penaeus vannamei*; Zm, *Zea mays*; Si, *Setaria italica*; Os, *Oryza sativa*; Ta, *Triticum aestivum*; Bd, *Brachypodium distachyon*; Ac, *Coffea arabica*; Ca, *Capsicum annuum*; Dc, *Daucus carota*; Gm, *Glycine max*; Rc, *Rosa chinensis*; At, *Arabidopsis thaliana*; Pp, *Physcomitrium patens*; Cr, *Chlamydomonas reinhardtii*. UniProt (https://www.uniprot.org) entries of the proteins are marked behind the species abbreviations. **b**, Tandem PHD sequence alignment for the corresponding species in A. Identical residues are highlighted in red. The dots indicate the critical residues responsible for interaction with H3K4me1 in the co-crystal structure of the Ehd3 tandem PHD–H3K4me1 complex. **c**, ITC results showing the binding affinities of homologous tandem PHD in *C. reinhardtii*, *P. patens*, *A. thaliana*, and *M. musculus* with H3K4me0 and methylated H3K4 peptides. NDB, no detectable binding.

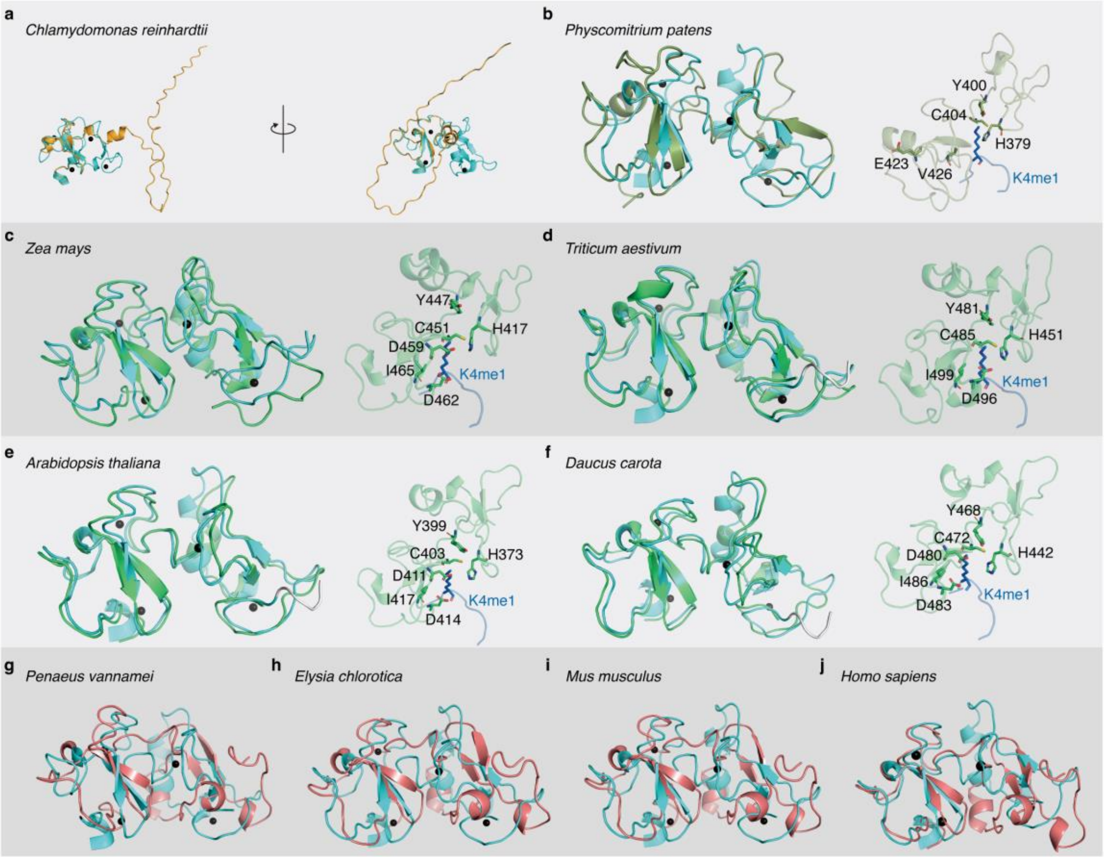


**Extended Data Fig. 9: Structural conservation of the tandem PHD finger domain in land plants.**

**a**–**j**, Structure superimposition of the tandem plant homeodomain (PHD) in A0A2K3D188 (a), A0A2K1L966 (b), A0A1D6LDK6 (c), A0A3B6RKR3 (d), Q1JPM3 (e), A0A162A0W2 (f), A0A3R7Q333 (g), A0A433TT05 (h), Q8BRH4 (KMT2C) (i), and O14686 (KMT2D) (j) with Ehd3. The structure of Ehd3 is colored in cyan. The protein codes represent the UniProt entries (https://www.uniprot.org).

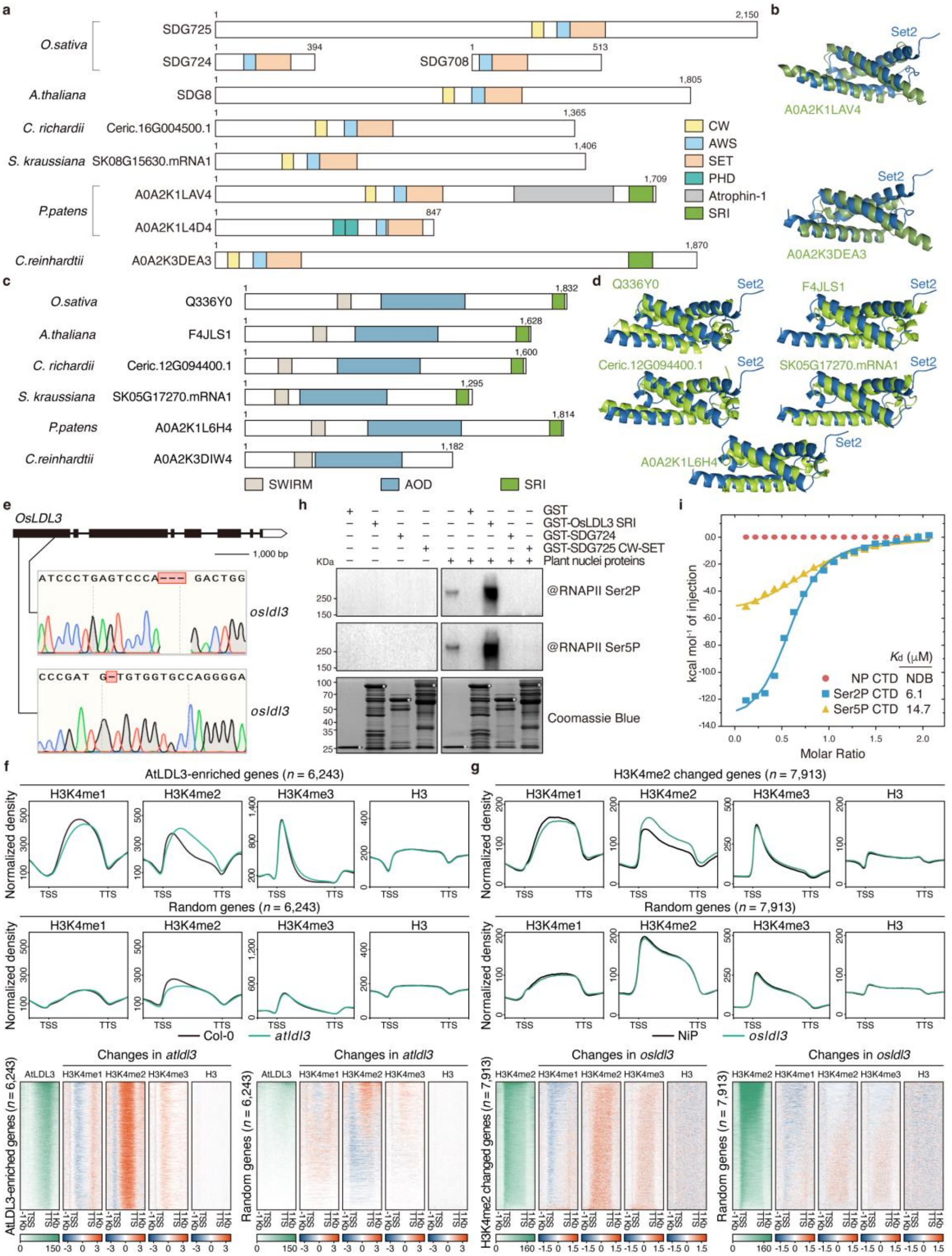


**Extended Data Fig. 10: Plant H3K4 demethylase LDL3 interacts with phosphorylated RNA polymerase II (RNAPII) via the Set2–Rpb1 interacting (SRI) domain, facilitating demethylation of H3K4me2 to H3K4me1.**

**a,** Schematic representation of H3K36 methyltransferases in *Chlamydomonas reinhardtii*, *Physcomitrium patens*, *Selaginella kraussiana*, *Ceratopteris richardii*, *Arabidopsis thaliana*, and *Oryza sativa*. **b,** Structure superimposition of the SRI domain in *Chlamydomonas reinhardtii* A0A2K3DEA3 and *Physcomitrium patens* A0A2K1LAV4 with *Saccharomyces cerevisiae* Set2 (PDB: 2a7o). **c,** Schematic representation of LDL3 homologous proteins in *Chlamydomonas reinhardtii*, *Physcomitrium patens*, *Selaginella kraussiana*, *Ceratopteris richardii*, *Arabidopsis thaliana*, and *Oryza sativa*. **d,** Structure superimposition of the SRI domain in *Physcomitrium patens* A0A2K1L6H4, *Selaginella kraussiana* SK05G17270.mRNA1, *Ceratopteris richardii* Ceric.12G094400.1, *Arabidopsis thaliana* F4JLS1, and *Oryza sativa* Q336Y0 with *Saccharomyces cerevisiae* Set2 (PDB: 2a7o). The protein codes represent the UniProt entries (https://www.uniprot.org), except for *Selaginella kraussiana* and *Ceratopteris richardii*, which are from https://sk.ccgg.fun/blast. **e,** Identification of rice *osldl3* mutants generated by CRISPR/Cas9 editing. The untranslated regions, exons, and introns of *OsLDL3* are indicated by blank rectangles, black rectangles, and black lines, respectively. **f**,**g**, Integrative genomic distributions and heatmaps showing the occupancies and changes of H3K4me1, H3K4me2, H3K4me3, and H3 in *Arabidopsis thaliana* wild-type Columbia-0 (Col-0) and the *atldl3* mutant (f) and *Oryza sativa* wild-type 'Nipponbare' (NiP) and the *osldl3* mutant (g). Genes enriched in AtLDL3 **(f) o**r with H3K4me2 changes **(g)** were analyzed. An equal number of random genes without AtLDL3 enrichment (f) or H3K4me2 changes (g) were used as negative controls. The plots and heatmaps present the region from 1 kb upstream of transcription start site (TSS) to 1 kb downstream of transcription termination site (TTS). **h**, Semi-*in vivo* pull-down showing the interaction between OsLDL3 and Ser-2- or Ser-5-phosphorylated RNA polymerase II (RNAPII). Total nuclear proteins extracted from NiP seedlings were precipitated using glutathione-*S*-transferase (GST)–OsLDL3 SRI, GST–SDG724, and GST–SDG725 CW-SET. GST served as a negative control. The products were analyzed by Coomassie Brilliant Blue staining and western blotting using antibodies against Ser-2- and Ser-5-phosphorylated RNAPII. The asterisk represents the corresponding recombinant protein. **i**, Isothermal titration calorimetry results showing the binding affinities of the SRI domain of

OsLDL3 with different modifications of the RNAPII C-terminal domain (CTD). NP, non-phosphorylated CTD peptide; Ser2P, Ser-2-phosphorylated CTD peptide; Ser5P, Ser-5-phosphorylated CTD peptide; NDB, no detectable binding.
